# *Moraxella catarrhalis* HemW is a Heme-binding Radical SAM Enzyme

**DOI:** 10.64898/2026.01.24.701380

**Authors:** Caitlin M. Padgett, Melissa M. Bollmeyer, Kaleb Boswinkle, Valérie de Crécy-Lagard, R. David Britt, Wen Zhu

## Abstract

*Moraxella catarrhalis* is an emerging human respiratory pathogen responsible for a significant proportion of childhood otitis media and exacerbations of chronic obstructive pulmonary disease. Recent transposon sequencing analysis identified *yggW* (renamed as *hemW*), an uncharacterized gene, as essential for *M. catarrhalis* growth under iron-limiting conditions, mimicking host-imposed nutritional immunity. HemW is annotated as a putative radical *S*-adenosylmethionine (SAM) enzyme and belongs to the HemN-like subfamily, but its biochemical properties remain unclear. Here, we report on the first experimental characterizations of *Mc*HemW. Our bioinformatic analysis confirmed its evolutionary relationship with the putative heme-binding radical SAM enzyme *Ec*HemW in *Escherichia coli*. Using biochemical and spectroscopic approaches, we demonstrate that *Mc*HemW contains a catalytically active [4Fe-4S] cluster and binds heme *in vitro*. These findings support a functional role for *Mc*HemW as a putative heme chaperone, highlighting it as a potential target for disrupting iron metabolism in this clinically important pathogen.

## Introduction

*Moraxella catarrhalis* is a bacterial pathogen associated with infections of the human upper and lower respiratory tract.^1,2^ Despite its clinical significance, our understanding of its metabolic vulnerabilities remains limited.^3,4^ This knowledge gap is particularly pressing given the rising prevalence of antibiotic resistance and the urgent need to identify novel targets for antimicrobial development.^5–7^ Using genomic array footprinting, along with targeted gene deletion and complementation, recent studies identified *yggW* (renamed *hemW*) as a previously uncharacterized gene essential for the growth of pathogenic *M. catarrhalis* BBH18 under iron-starvation conditions.^8^ Essential genes required for bacterial growth under iron-limiting conditions are thought to reflect the physiological stresses encountered by pathogens within the human host.^9–11^ The conserved requirement for *hemW* across multiple *M. catarrhalis* isolates suggests that it plays a critical role in their pathogenicity.^8^

The protein encoded by *M. catarrhalis hemW* is annotated as a putative radical *S*-adenosylmethionine (SAM) enzyme, HemW. With over 800,000 members, the radical SAM superfamily has emerged as one of the largest and most functionally diverse superfamilies, carrying out essential biological functions across all kingdoms of life.^12–15^ Most members of this superfamily share a conserved CX_3_CX_2_C motif in the α_6_β_6_ half-TIM barrel used to coordinate a [4Fe-4S] cluster.^16–18^ This [4Fe-4S] cluster mediates the reductive cleavage of SAM, initiating a cascade of radical-based reactions that enable a wide range of distinct transformations of a second substrate, typically with high regio-and stereoselective.^19–22^ Chemical versatility and broad substrate scope of these enzymes arise from auxiliary domains appended to the conserved half-TIM barrel core (Figure 1a).^12,23–35^ Functional annotation of many radical SAM subfamilies is often hindered by poorly characterized auxiliary domains.^36–38^ An under characterized subgroup of the radical SAM superfamily is the oxygen-independent coproporphyrinogen III oxidase-like enzymes, sometimes referred to as HemN-like radical SAM enzymes.^39–41^ These enzymes are proposed to use two equivalents of SAM to catalyze C-C bond-forming or cleaving reactions on diverse substrates, such as peptides and polyketides.^42–46^ The only X-ray crystal structure available for this subfamily,^47^ HemN, reveals two distinct SAM-binding sites and supports a proposed mechanism in which one SAM molecule, coordinated to the [4Fe-4S] cluster, generates the 5’-deoxyadenosyl (5-dAdo) radical, while a second SAM generates a methylene radical to initiate oxidative decarboxylation of protoporphyrin IX (Figure 1b).^48^ Several members of this subfamily, such as HemN, ChuW, and HutW, are involved in heme biosynthesis and degradation pathways (Figure 1b),^49–53^ however, the vast majority of enzymes within this subfamily await functional elucidation.

**Figure 1.**
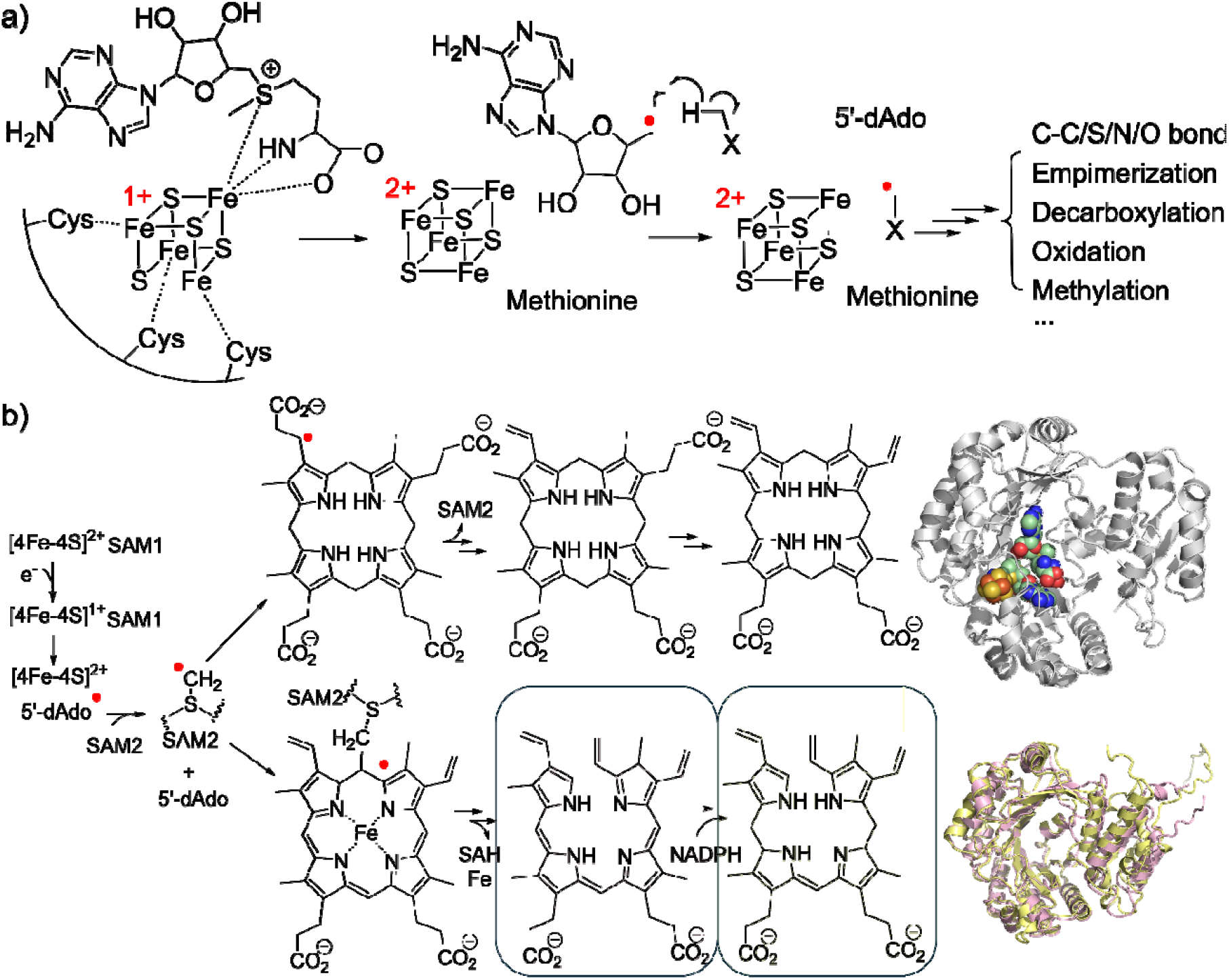
Overview of radical SAM enzymes involved in heme biosynthesis and degradation. a) Canonical radical SAM enzyme uses a [4Fe-4S] cluster to reductively cleave SAM to initiate a cascade of reactions. b) Left: Mechanisms of HemN (top) and ChuW/HutW (bottom). Both pathways employ a canonical [4Fe-4S] cluster that mediates reductive cleavage of the first SAM molecule to generate the 5’-dAdo radical, followed by use of a second SAM for C-C bond cleavage. The pink box indicates the tetrapyrrole product generated by ChuW, and the yellow box shows the corresponding product formed by HutW. Top right: X-ray crystal structure of HemW (PDB:1OLT) with the [4Fe-4S] cluster (Fe, orange; S, yellow spheres) and bound SAM (C, pale green; N, blue; O, red). Bottom right: AlphaFold4-predicted structural models of *E. coli* O157:H7 ChuW (pink) and *Vibrio cholerae* HutW (yellow), highlighting overall fold similarities among these radical SAM enzymes.

In this study, we heterologously expressed and purified *M. catarrhalis* HemW. Using a combination of bioinformatics, biochemical, and spectroscopic approaches, we demonstrate that *Mc*HemW is capable of binding heme and harbors a catalytically active [4Fe-4S] cluster. We also observed that heme binding enhances the SAM cleavage activity. These findings suggest a putative role for *Mc*HemW as a catalytically active radical SAM enzyme with heme-binding capability, providing a fundamental mechanistic framework for future efforts to uncover the relevant biological pathways and their role in the pathogenesis of *M. catarrhalis*.

## Results

### Sequence similarity network (SSN) analysis identifies *Mc*HemW as a putative heme chaperone

The sequence of *Mc*HemW contains a characteristic CX_3_CX_2_C motif, indicative of a putative radical SAM enzyme. Sequence similarity analysis using RadicalSAM.org^36^ revealed that *Mc*HemW belongs to the oxygen-independent coproporphyrinogen III oxidase-like enzymes, which are found across bacteria, archaea, and eukaryotes (Figure S1). Only a few members of this subfamily have been experimentally characterized, including HemN^48,49,54–58^ and HemZ^59,60^, which have been identified to participate in heme biosynthesis, while others, such as ChuW^50^ and HutW^51,54^, are involved in heme degradation pathways. Heme-degradation radical SAM enzymes are proposed to share a similar radical-initiation step, with divergent substrate modifications (Figure 1b).^50,51,54^ The location of the protoporphyrin-binding pocket has been proposed to be at the auxiliary domain based on computational modeling. When we systematically increased sequence alignment stringency until previously characterized enzymes segregated into distinct functional clusters, at the alignment score 110 threshold, *Mc*HemW grouped within the same cluster as *Escherichia coli* HemW, *Ec*HemW, a putative heme chaperone. (Figure 2a and Figure S2).^61^ We also examined the phylogenetic relationships among previously characterized radical SAM enzymes involved in heme-related pathways (Table S1). Our analysis revealed a clear separation between HemN and HemW enzymes (Figure□2b). Within the HemW clade, *Lactococcus lactis* HemW^62^, *Ll*HemW, forms a distinct branch, while *Mc*HemW clusters closely with *Ec*HemW^61^, sharing 43% sequence identity. This is consistent with our SSN analysis, which shows that *Mc*HemW shares high sequence similarity with *Ec*HemW (Figure S2).

**Figure 2.**
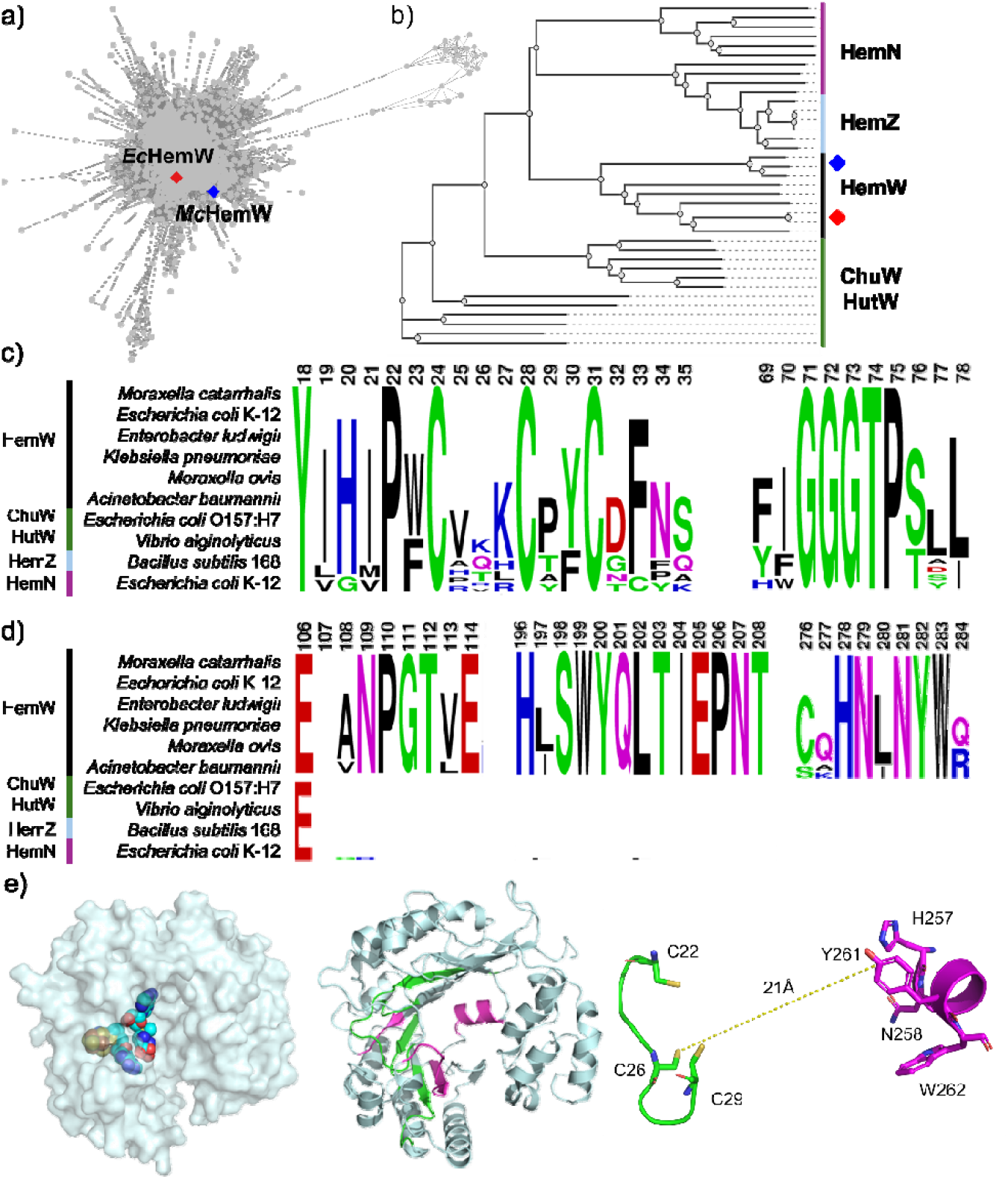
Bioinformatic analysis of *Mc*HemW. a) Sequence similarity network (SSN) identifies *Mc*HemW (blue diamond) clustering with *Ec*HemW (red diamond). b) Maximum-likelihood phylogenetic tree showing the evolutionary relationships among HemW, ChuW, HutW, HemZ, and HemN homologs. HemW cluster distinctly from HemZ/HemN clades. Colored diamonds are the same as in panel a. c) Sequence logo representations of conserved SAM-binding motifs across HemW, HemZ, ChuW, and HutW homologs, highlighting residues that define the shared radical SAM catalytic core. d) Sequence conservation patterns unique to HemW homologs, showing regions that are highly conserved within HemW but absent or divergent in other heme-binding radical SAM enzymes. e) Left: Surface representation of the AlphaFold4 model of *Mc*HemW, with modeled [4Fe–4S] cluster (yellow and orange spheres) and bound SAM (C, cyan; N, blue; O, red). Middle: Structural mapping of HemW-specific conserved regions (magenta) compared with features shared across other radical SAM enzymes (green), illustrating elements that distinguish HemW within the broader subfamily. Right: Estimated distance between CX_3_CX_2_C motif and the conserved HNX_2_YW motif residues of the AlphaFold4 model of *Mc*HemW.

To better define the sequence features that distinguish *Mc*HemW from neighboring subclusters, we performed multiple sequence alignment analysis on HemW and other radical SAM enzymes involved in the heme biosynthesis and degradation pathways (Figure S3). We observe that *Mc*HemW contains C22-C26-C29 and G71-G72-G73-T74-P75 motifs, which are required for [4Fe-4S] cluster and SAM binding (Figure 2c). The alignments further indicate that the principal distinction between HemN and HemW is associated with the residues involved in coordinating the second SAM-binding site (Figure□2d). Mapping these regions onto the AlphaFold-predicted *Mc*HemW structural model (Figure□2e) indicates that canonical radical SAM features, such as the [4Fe-4S] cluster-binding motif and the surrounding SAM-binding pocket, are likely conserved in *Mc*HemW. In contrast, the distal cavity adjacent to the active-site residues appears to be unique to HemW homologs and is not observed in ChuW, HutW, HemZ, or HemN.

### *Mc*HemW contains an iron-sulfur cluster

We expressed and purified His-tagged *Mc*HemW in *E. coli* BL21-Gold (DE3) under anaerobic conditions. Intact protein mass spectrometry confirmed a mass of 46,137 Da, consistent with the expected sequence (Figure S4). The purified protein is brown in color and migrates as a single band on SDS-PAGE (Figure 3a). The iron and sulfur content of the as-purified sample was 3.1 ± 0.7 and 3.4 ± 0.5 per monomer, respectively. Chemical reconstitution gave 6.1 ± 0.6 Fe and 5.9 ± 0.9 S per monomer, indicating the presence of at least one iron-sulfur cluster and likely binding of adventitious iron and sulfur. In the UV-visible spectrum, we observed the broad absorption feature at 410 nm of *Mc*HemW, and the addition of sodium dithionite (DTH) abolished this feature (Figure 3b). To test if the iron-sulfur cluster is coordinated by the conserved CX_3_CX_2_C motif within the half-TIM barrel, we generated a double-cysteine knockout mutant, C22A/C29A, to disrupt the putative radical SAM (RS) cluster-binding site. This mutant, designated ΔRS, was purified to homogeneity at a level comparable to that of the wild-type (WT) protein but lacked the characteristic brown color (Figure 3a), showed no absorbance at 410 nm and contained no detectable iron (Figure 3b). We conclude that *Mc*HemW coordinates an iron-sulfur cluster via the C22-C26-C29 motif.

**Figure 3.**
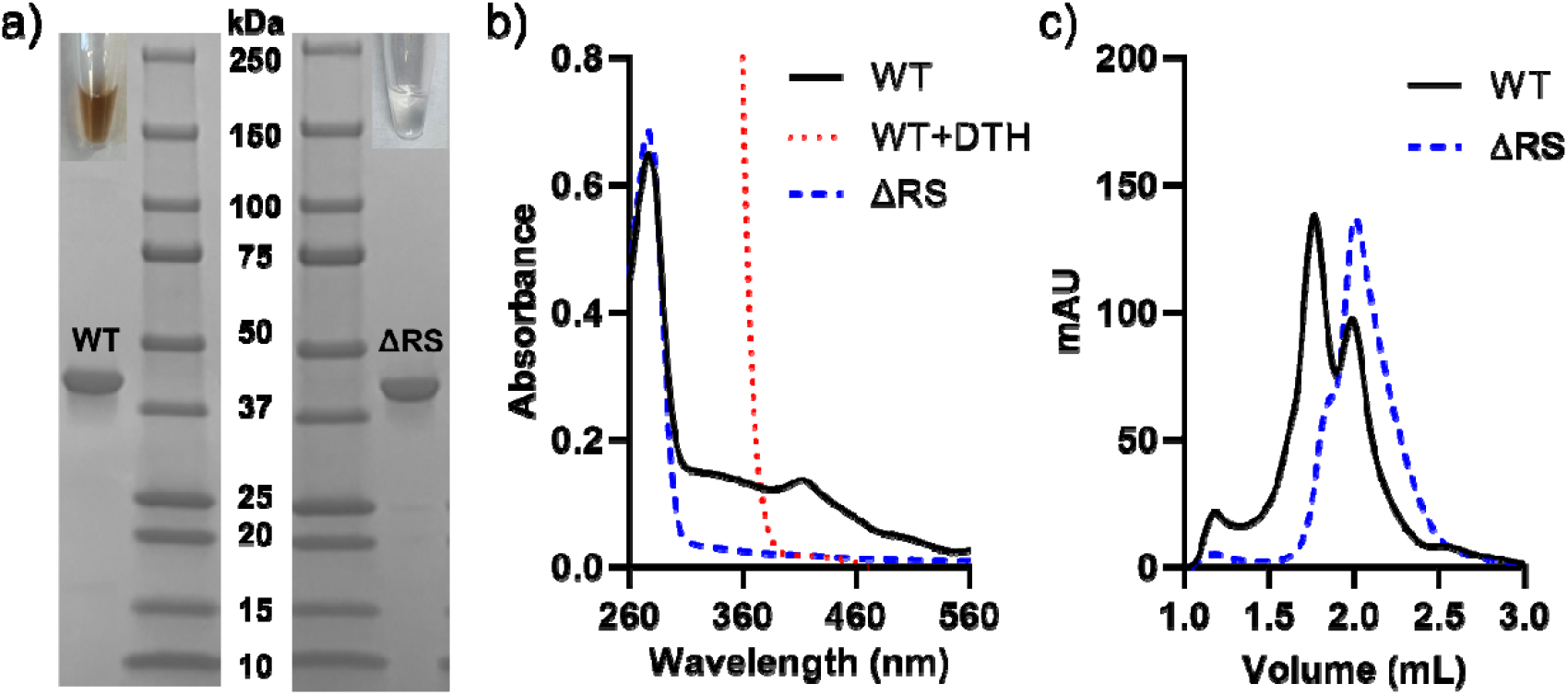
The CX_3_CX_2_C motif in *Mc*HemW coordinates the iron-sulfur cluster, and the cofactor is required to maintain the enzyme’s dimeric state. a) SDS-PAGE analysis of purified WT *Mc*HemW and the ΔRS variant lacking the iron-sulfur cluster binding motif. Insets show the corresponding protein solutions after purification. WT HemW exhibits the characteristic brown coloration associated with [4Fe-4S] cluster, whereas ΔRS is colorless, consistent with the absence of cofactors. Both proteins are expressed and purified to comparable levels. b) UV-vis spectra of WT (black), ΔRS HemW (blue dashed), and WT with the addition of DTH (dotted red). WT *Mc*HemW displays distinct visible absorption features indicative of iron-sulfur cluster binding. These features are absent in ΔRS and reduced WT samples c) Size-exclusion chromatography shows that WT HemW (black) exists as a mixture of dimer and monomer in solution, whereas the ΔRS variant (blue dashed) is predominantly monomeric.

Native-PAGE analysis revealed that chemically reconstituted *Mc*HemW exists in both monomeric and dimeric forms in solution (Figure S7). To further characterize its oligomeric state, we performed analytical size-exclusion chromatography (SEC) under anaerobic conditions (Figure 3c). Two distinct peaks corresponding to ∼46 kDa and ∼100 kDa confirmed the presence of both monomer and dimer species. Increasing protein concentration does not significantly shift the relative abundance of the dimer (Figure S4). In contrast, the ΔRS variant, which lacks the iron-sulfur cluster, predominantly existed as a monomer at the same concentration (Figure 3c). These findings are consistent with previous observations for *Ec*HemW, in which the intact iron-sulfur cluster binding site promotes protein dimerization.^57^

### EPR characterization of [4Fe-4S] cluster coordination

The continuous-wave (CW) X-band EPR spectrum of chemically reconstituted *Mc*HemW contained a signal at *g* = 4.3 attributed to adventitious Fe (III) and a sharp signal at *g* = 2.01 that we assign to a [3Fe-4S]^+^ cluster, which could be the result of incomplete cofactor incorporation (Figure 4a). Upon the addition of DTH, a new S = 1/2 signal appeared with a *g*-tensor of [1.99, 1.92, 1.82] (Table 1). The *g* = 2.01 feature remained after the addition of DTH, indicating that the [3Fe-4S]^+^ cluster was not fully reduced. Both signals were absent in the spectra of the ΔRS variant and can, therefore, be attributed to the RS cluster. A simulation of the reduced *Mc*HemW spectrum indicates that the signal with *g*_max_ = 1.99 is the dominant species, accounting for approximately 97% of the total spins detected (Figure 4b). The low *g*_max_ value of 1.99 is more consistent with a [2Fe-2S]^+^ cluster, whereas [4Fe-4S]^+^ clusters typically have *g*_max_ values greater than *g*_e_.^63,64^ However, increasing the temperature completely diminished this signal, leaving only the signal corresponding to the [3Fe-4S]^+^ cluster. This fast-relaxing behavior is more consistent with [4Fe-4S]^+^ clusters, while [2Fe-2S]^+^ clusters tend to relax more slowly and retain signal even at 60 K.^65^ The low value of *g*_max_ and fast relaxing behavior of this signal make the assignment of the cluster as either a [2Fe-2S]^+^ or [4Fe-4S]^+^ species ambiguous.

**Table 1.** Simulation parameters used for the reduced *Mc*HemW EPR spectra.

| Component | *g*-tensor | weight %<br>reduced *McHemW* | weight %<br>reduced *McHemW* + CN |
| --- | --- | --- | --- |
| [4Fe-4S]<sup>+</sup> | 1.99, 1.92, 1.83 | 97 | 61 |
| [3Fe-4S]<sup>+</sup> | 2.01 | 3 | 6 |
| CN-[4Fe-4S]<sup>+</sup> | 2.07, 1.95, 1.91 | 0 | 33 |

**Figure 4.**
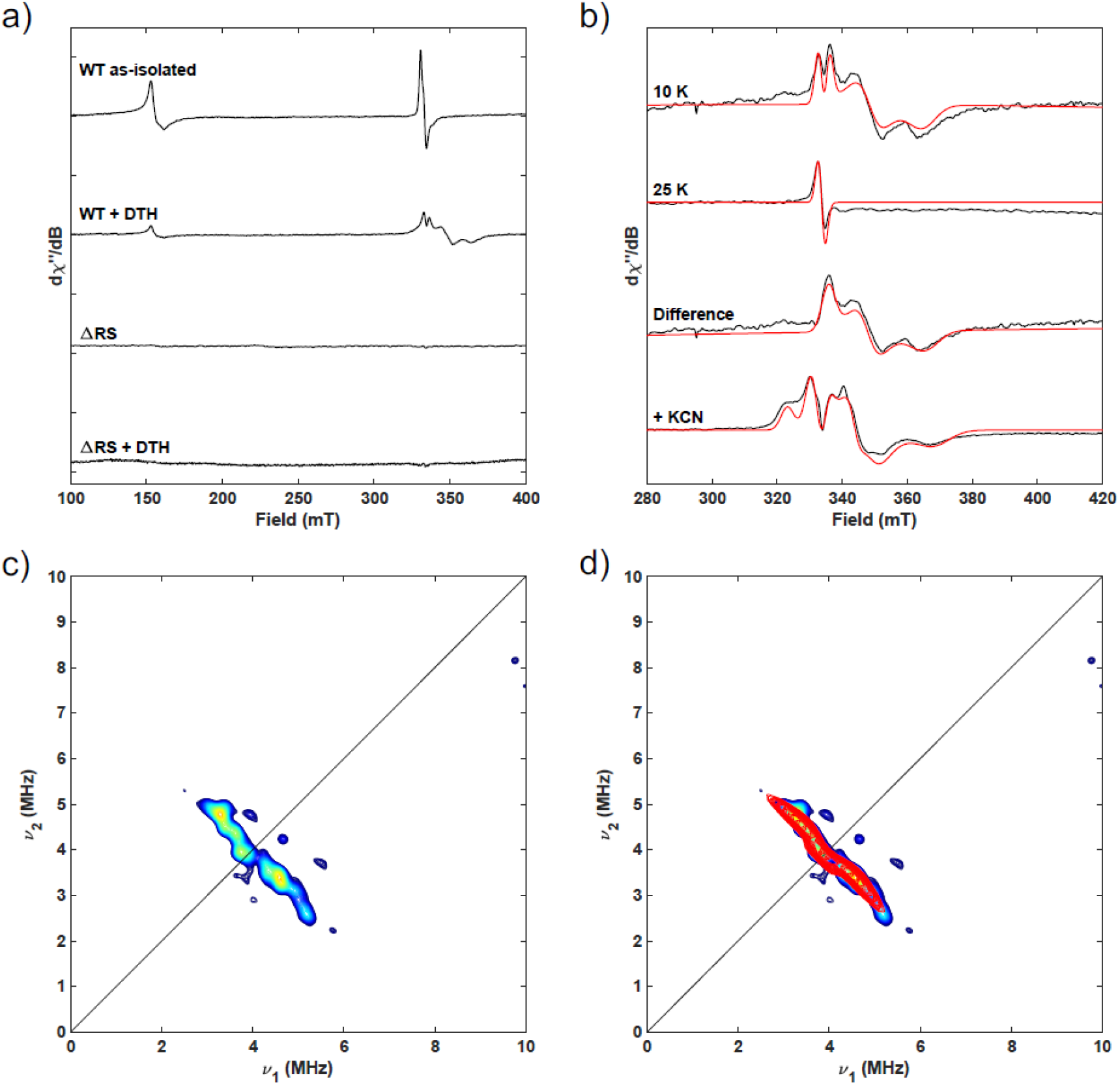
X-band EPR spectra of *Mc*HemW samples. a) CW spectra comparing before and after dithionite (DTH) reduced *Mc*HemW with the ΔRS variant. b) Narrow scans of the S =1/2 signals for reduced *Mc*HemW recorded at 10 K, 25 K, the difference spectrum (10 K-25 K), and the CN-bound *Mc*HemW spectrum recorded at 10 K. Experimental spectra are shown in black and spectral simulations are shown in red. C) HYSCORE spectrum collected at g = 1.94 for DTH-reduced *Mc*HemW with the addition of 100 equivalents of K^13^CN. D) The simulated HYSCORE spectrum is shown in red overlayed with the experimental spectrum. Simulation parameters: *A*^13^C = [−5.0, −4.5, 1.9] MHz and Euler angle = [0 20 0]°. Experimental parameters: for CW spectra, frequency = 9.4 GHz, temperature = 10 K (unless noted), power = 0.2 mW, modulation amplitude = 5 G. For HYSCORE spectra, temperature = 10 K, *t*_π/2_ = 12 ns, *t*_π_ = 24 ns, frequency = 9.195 GHz, field = 338.6 mT, and τ = 140 ns. The time increment in both dimensions was 24 ns with 180 steps.

To establish whether the reduced iron-sulfur cluster signal could be assigned to a [4Fe-4S]^+^ RS cluster, we used ^13^C-labeled cyanide as a probe. Cyanide can bind to the unique Fe of an RS cluster, but cannot bind to a cluster fully coordinated by cysteine residues in a [2Fe-2S] cluster. When reduced *Mc*HemW was incubated with 100 equivalents of K^13^CN, a new species with *g* = [2.07, 1.95, 1.91] appeared (Figure 4b). This new rhombic *g*-tensor is similar to those of other ^13^CN-bound RS clusters, including PqqE, IspH, HydG, and *P. furiosus* ferredoxin.^66–70^ While this signal is consistent with a ^13^CN-bound [4Fe-4S]^+^ RS species, some of the unbound [4Fe-4S]^+^ cluster remained, indicating that there was incomplete binding. Further characterization of ^13^CN-treated HemW using X-band hyperfine sublevel correlation (HYSCORE) spectroscopy acquired at *g* = 1.94 revealed one set of cross-peaks centered at the ^13^C Larmor frequency (Figure 4c).

These cross-peaks were well simulated using a pseudoaxial hyperfine coupling tensor *A*^13^C = [−5.0, −4.5, 1.9] MHz and Euler angle = [0, 20, 0]° (Figure 4d). This hyperfine tensor is similar to those observed for other ^13^CN-ligated [4Fe-4S]^+^ clusters and provides additional support for the presence of a typical [4Fe-4S]-loaded RS cluster.^66–70^

### Heme binds to *Mc*HemW non-covalently

A previous study of *Ec*HemW showed that the protein covalently binds heme in a 1:1 stoichiometry *in vitro*. To explore the nature of heme binding in *Mc*HemW, we reconstituted *Mc*HemW with heme at a 1:1 molar ratio and analyzed the complex by UV-vis spectroscopy to assess its heme-binding capability (Figure 5a). Heme in aqueous solution exhibited a broad absorption band at 350-385 nm, with a weak absorption peak at 600 nm. The heme-reconstituted *Mc*HemW displayed a sharp Soret band at 411 nm, suggesting that heme is bound in protein at an oxidized Fe (III) state within an electronic environment distinct from that of free heme. In contrast, with the ΔRS variant, the heme spectrum showed only weak, red-shifted features at 360-390 nm, with no discernible Soret band signals. This feature was absent in the sample in which HemW was replaced with bovine serum albumin (BSA). The addition of DTH to WT *Mc*HemW results in a red shift of the Soret band to 420 nm, accompanied by an increase in intensity, and the appearance of the distinct Q-bands at 532 nm and 562 nm (Figure 5b). The appearance of the 560 nm Q-band likely reflects ligand-induced changes, possibly due to the reduction to Fe (II) or stabilization of a defined spin state. When heme was titrated into a fixed concentration of *Mc*HemW, binding was monitored by changes in the Soret absorbance (Figure 5c). Analysis of the 411 nm intensity, normalized against the free-heme background at the same wavelength, showed a progressive increase in the Soret signal with increasing heme concentrations (Figure S5). This indicates that the elevated absorbance at 411 nm arises from the formation of the *Mc*HemW-heme complex rather than from the higher background signal of free heme. Unfortunately, the high concentration of heme saturated the UV-vis detection, therefore, limiting our ability to quantify the binding affinity. We estimated, however, that the binding affinity is in the micromolar range, consistent with the previously observed HemW homolog in *Campylobacter jejuni*.^71^ Our UV-Vis spectra indicate that *Mc*HemW binds heme, and that this binding benefited from an intact [4Fe-4S] cluster or the integrity of its binding site.

**Figure 5.**
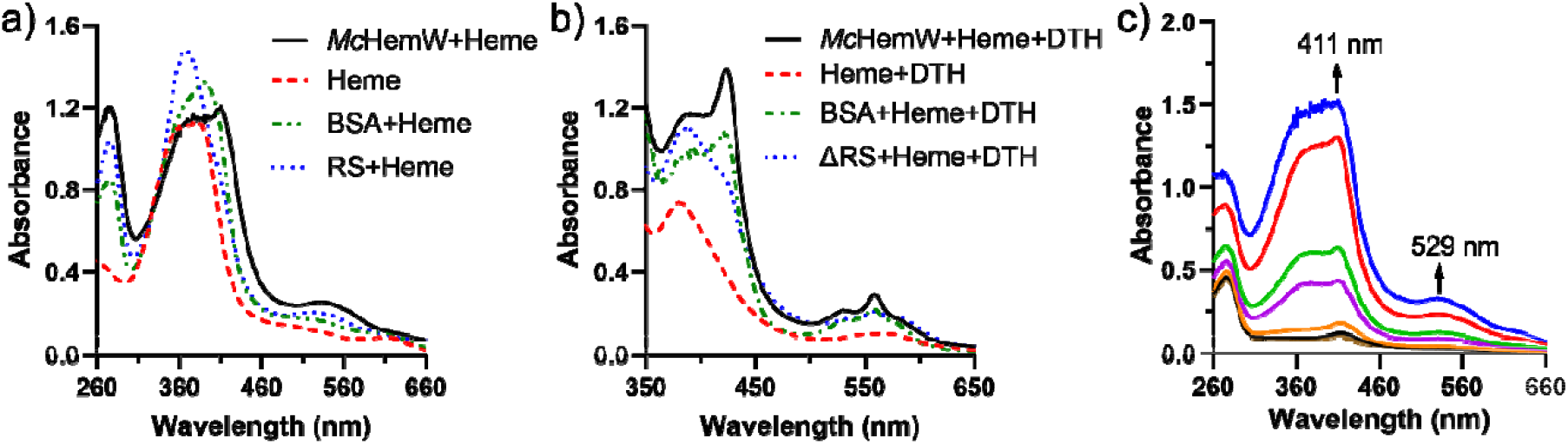
Heme-binding of *Mc*HemW and the ΔRS variant reveals loss of Fe-S and heme features upon disruption of the iron-sulfur cluster. a) UV-vis spectra of WT *Mc*HemW show a broad visible-region absorbance from free heme and a pronounced Soret band corresponding to bound heme. These heme-associated features are absent in both the ΔRS variant and the BSA control. b) UV-vis spectra of WT *Mc*HemW under DTH-reduced conditions show a pronounced Soret band corresponding to bound heme. These heme-associated features are absent in both the ΔRS variant and the BSA control. c) UV-vis spectra of *Mc*HemW bound to increasing concentrations of heme reveal a dose-dependent enhancement of the Soret band near ∼400 nm and additional visible-region features, consistent with heme association. 5 μM WT *Mc*HemW with increasing concentrations of heme 0 μM (brown), 0.5 μM (black), 1 μM (orange), 3.5 μM (purple), 5 μM (green), 10 μM (red), and 15 μM (blue).

We next investigated whether the dimer and monomer of *Mc*HemW are both capable of binding heme. To this end, we performed native-PAGE on heme-reconstituted WT and the ΔRS variant, followed by dual staining with 3,3′,5,5′-tetramethylbenzidine (TMBZ) for heme detection and Coomassie Blue for total protein analysis (Figure S7). We observed both the monomer and the dimer of WT enzyme in the native-PAGE gel, consistent with our SEC analysis. When the same gel was treated with TMBZ, both dimeric and monomeric forms of WT *Mc*HemW stained with TMBZ at high protein concentration, indicating that both forms are competent for heme binding. In contrast, the ΔRS variant contains a monomeric form on native-PAGE when stained with Coomassie Blue. TMBZ staining of the ΔRS variant also shows a much weaker band than the same concentration in the WT sample at the size corresponding to the dimeric form of the ΔRS variant on native-PAGE. When we repeated the dual staining experiment on SDS-PAGE gels using heme-reconstituted WT and the ΔRS variant, no heme staining was observed in the ΔRS variant, despite the protein band being readily visible on the Coomassie stain (Figure S7). We observed weak heme binding only at high protein concentrations (50 μM) for WT, consistent with similar interactions previously reported for *Ec*HemW.^61^ We suspected that the heme staining observed at high protein concentrations arises from incomplete denaturation in an overloaded SDS-PAGE. To confirm this hypothesis, we performed intact protein mass spectrometry on heme-reconstituted WT *Mc*HemW. Upon incubation with heme, no shift in protein mass was detected, and only free heme was detected at the expected mass of 616 Da, confirming that heme does not form a covalent adduct with *Mc*HemW (Figure S6). As the ΔRS variant disrupts [4Fe-4S] cluster coordination and impairs dimerization, the difference between WT and ΔRS suggests that the heme-binding favors the enzyme containing an intact [4Fe-4S] cluster or its binding motif, likely due to the influence on the local conformation of the heme coordination site.

### EPR characterization of heme binding

When heme was added to *Mc*HemW, the EPR spectrum of the oxidized sample featured a new low-spin heme signal in addition to the [3Fe-4S]^+^ signal near *g* = 2 (Figure 6). This low-spin heme signal is distinct from the axial S = 5/2 signal of heme and is indicative of heme binding to the enzyme.^72,73^ The spectrum of heme-bound *Mc*HemW was best fitted with two S = 1/2 species with *g* = [2.45, 2.28, 1.90] and *g* = [2.52, 2.27, 1.87] in a 2:1 ratio. Differences in heme conformation or axial ligation could account for the two distinct sets of *g* values. The observed heterogeneity in heme binding likely reflects the mixture of dimeric and monomeric protein populations. In the heme-reconstituted ΔRS variant, we observed two closely related species, with the second heme species exhibiting slightly increased rhombicity, characterized by *g* values of [2.53, 2.28, 1.86]. Since we observe a minor dimeric form in the ΔRS variant by SEC and native-PAGE, the more rhombic species (*g* = [2.53, 2.28, 1.86]) in the ΔRS variant of *Mc*HemW likely represent the heme bound to the monomeric form of the ΔRS variant, which has an altered binding conformation. This indicates that binding of the [4Fe-4S] cluster is important for the overall fold of the enzyme and its interaction with heme.

**Figure 6.**
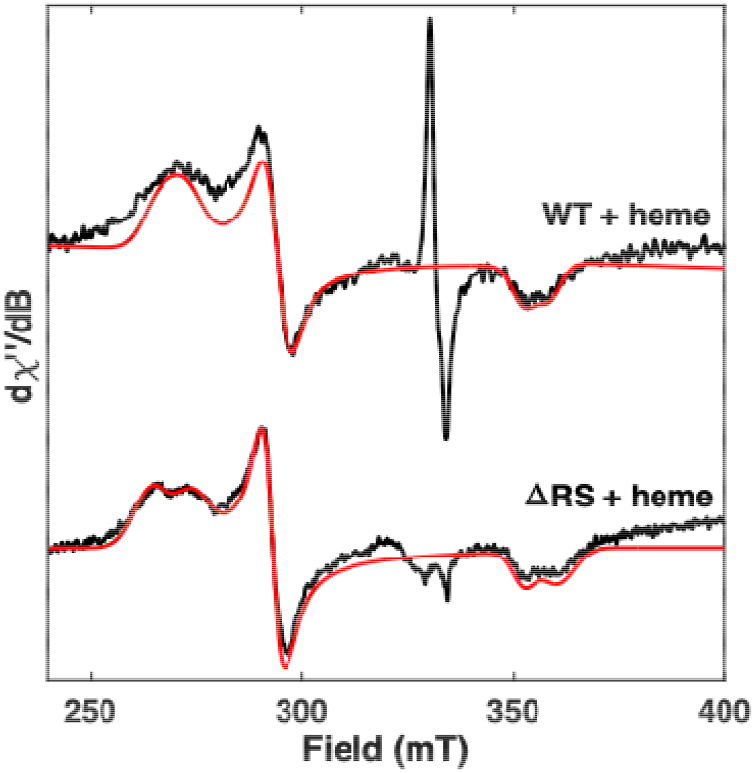
CW X-band EPR spectra of *Mc*HemW with heme or the ΔRS variant with heme. Experimental spectra are shown in black and spectral simulations are shown in red. Experimental parameters: Frequency = 9.4 GHz, temperature = 10 K, power = 0.2 mW, modulation amplitude = 5 G. Simulations: top) *g* = [2.45, 2.28, 1.90] and *g* = [2.52, 2.27, 1.87]. bottom) *g* = [2.45, 2.28, 1.90] and *g* = [2.53, 2.28, 1.86].

### Catalytic competence of the *Mc*HemW

Our sequence alignment and spectroscopic analyses suggest that *Mc*HemW uses the conserved CX_3_CX_2_C motif to coordinate a [4Fe-4S] cluster with an open Fe site for SAM binding, suggesting potential catalytic competence. When *Mc*HemW was incubated with SAM in the presence of DTH, HPLC analysis revealed new peaks eluting at the exact retention times of the commercial standards for 5’-dAdo and *S*-adenosylhomocysteine (SAH) (Figure 7a). LC-MS supports the identities of these products, which have 252.1 m/z and 385.1 m/z, respectively (Figure 7b). Omission of DTH, SAM, or *Mc*HemW abolished the formation of both products. A time-dependent activity assay showed that 5’-dAdo and SAH were formed with an initial rate of 0.26 ± 0.09 and 0.14 ± 0.05 μM/hr, respectively (Figure 7c). Intriguingly, when the activity assays were repeated using heme-reconstituted *Mc*HemW, the same product peaks were observed, but their intensities increased, resulting in 4.6-fold increases in the initial rates of 5’-dAdo and SAH formation (Figure 7c). The *Mc*HemW sample without chemical reconstitution showed a similar trend in our time-course measurement, with a slightly slower initial rate (Figure S8). Given that SAM and DTH are present in large excess, this low initial rate and low product conversion suggest that the reaction likely proceeds predominantly in a single-turnover manner. The amount of SAH formed is substoichiometric relative to 5’-dAdo, likely reflecting a shunt pathway operating during the reaction.^48,74^ However, the acceptor of the methyl group in this reaction is unclear because no heme-methylation or heme-degradation product was observed in our LC-MS analysis. Nonetheless, these results demonstrate that the RS [4Fe-4S] cluster of *Mc*HemW is catalytically competent to initiate radical chemistry, albeit much less efficiently than other radical SAM enzymes, such as *Bs*QueE.^75^ This observation is consistent with the basal activity observed for *Ec*HemW as reported previously. However, the presence of heme enhances SAM cleavage activity at the radical SAM site of *Mc*HemW, further implying the crosstalk between the radical SAM active site and the heme binding site.

**Figure 7.**
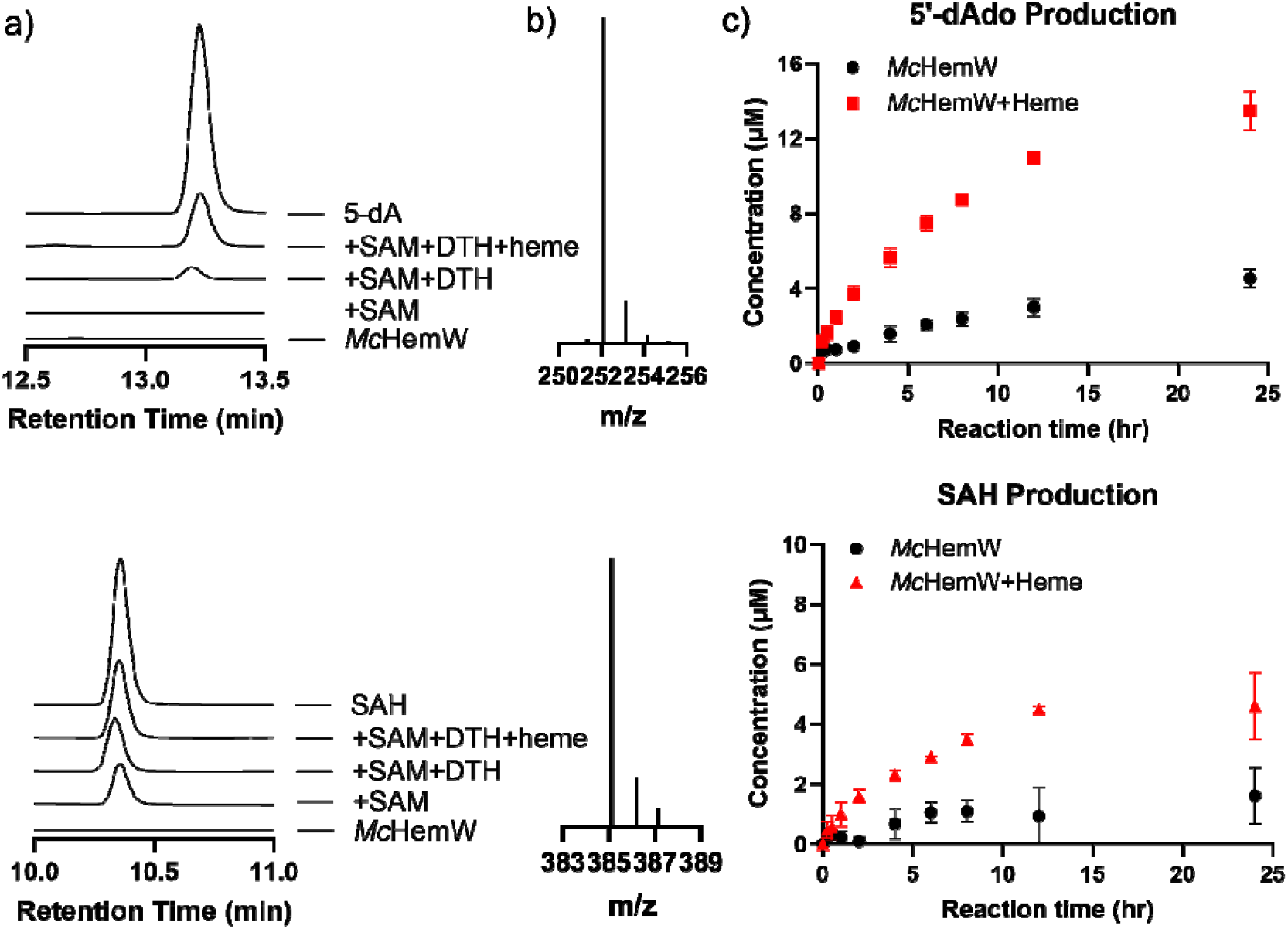
Activity assay of *Mc*HemW measured by HPLC and LC-MS. a) HPLC analysis identifies the production of 5’-dAdo and SAH by chemically reconstituted *Mc*HemW under different reaction conditions, including the presence or absence of DTH, heme, and SAM. b) LC-MS confirms the formation of both 5’-dAdo and SAH in the WT reactions. c) Time-course analysis of chemically reconstituted *Mc*HemW-catalyzed reactions shows that heme significantly enhances the production of 5’-dAdo and SAH. Error bars represent the standard deviation from two independent experiments.

## Discussion

*M. catarrhalis* is the third most common cause of childhood otitis media, or ear infections, accounting for 15-20% of cases.^76,77^ *M. catarrhalis* also causes approximately 10% of exacerbations of chronic obstructive pulmonary disease in adults, resulting in approximately 2-4 million episodes annually.^2,78^ Understanding the roles of *M. catarrhalis* genes is crucial for elucidating how *M. catarrhalis* survives within the host.

One of the key host defense strategies against bacterial pathogens is nutritional immunity, in which the host limits the availability of essential nutrients, such as iron, to restrict microbial growth and proliferation.^79–81^ Genes that support bacterial pathogen growth under iron-limited conditions have attracted particular attention in recent years due to their importance during infection.^82,83^ *M. catarrhalis* has multiple systems to overcome host-imposed iron restrictions, including heme uptake pathways, transferrin- and lactoferrin-binding proteins, and CopB-dependent transporters.^83–85^ The *hemW* gene is one of the essential genes for *M. catarrhalis* survival in this environment.^8^ In *E. coli* BW25113, deletion of *hemW* produces only a subtle phenotype under laboratory growth conditions.^61^ The HemW homolog in *Mycobacterium tuberculosis* is nonessential for *in vitro* growth on nutrient-rich media but is required for growth in the C57BL/6J mouse spleen.^86^ Differences across *in vitro* and *in vivo* conditions suggest that HemW-mediated processes may contribute to pathogen survival and virulence under host-relevant conditions.

Our bioinformatic analysis shows that *Mc*HemW is clearly separated from the previously characterized HemN homologs and constitutes a distinct subclass. In *M. catarrhalis*, the *Mc*HemW operon contains two additional genes, annotated as ATP-dependent RNA helicase RhIB and a hypothetical protein that does not appear to contribute directly to the iron-deficiency phenotype, suggesting a possible regulatory role (Figure S9).^8,87^ Similarly, the *Ec*HemW operon contains predicted regulatory genes hypothesized to participate in stress-response pathways that remodel gene expression under adverse conditions (Figure S9).^88^ Further comparative genomics analyses reveal two types of gene patterns of neighborhood conservation surrounding *hemW* (Figure S9). In α- and γ-proteobacteria, *hemW* frequently co-occurs with transcriptional activator *rdgB* and genes involved in nucleic acid-related metabolism, including *rph* (ribonuclease PH) (Figure S9).^89,90^ The functional implications of this association remain unclear, but it may hint at an unrecognized link between HemW and pathways that sense or respond to nucleic acids. A second, highly conserved pattern appears across many Gram-positive lineages and a subset of Gram-negative organisms, in which *hemW* is located adjacent to *hrcA* and canonical heat-shock responsive genes, such as *grpE, dnaK*, and *dnaJ*.^91–93^ In *Bacillus subtilis*, this linkage is experimentally validated, with *hemW* residing within a single operon that includes *lepA, hrcA, grpE*, and *dnaK*.^94^ The conservation of these genetic contexts suggests that HemW may interface with cellular stress pathways more broadly than previously appreciated. In particular, co-regulation with heat-shock genes could reflect a requirement for HemW function during proteotoxic or oxidative stress, consistent with its proposed role as a heme chaperone whose activity must remain robust.^61,95^ Although the mechanistic basis of these associations remains speculative, the genomic organization provides a compelling framework for future studies aimed at defining how HemW integrates into the bacterial stress response and cofactor trafficking networks.

We have shown that *Mc*HemW harbors an RS [4Fe-4S] cluster, which can be readily reduced by DTH. This behavior contrasts with previous reports on *Ec*HemW, in which the cluster could not be reduced by multiple electron donors, including sodium dithionite and titanium(III) citrate, thereby precluding successful EPR characterization, although the reduction potential determined by cyclic voltammetry matches that of a typical RS [4Fe-4S] cluster.^61^ The reported EPR study of HemN employed photochemical reduction mediated by excess sodium oxalate and 5-deazaflavin, a reductant system stronger than DTH.^49^ In comparison, EPR analyses of HutW, a heme-degrading enzyme, show that the enzyme can be efficiently reduced using DTH.^51,54^ We have also demonstrated that the oligomerization state of *Mc*HemW is dependent on the integrity of the [4Fe-4S] cluster binding site. Loss of the [4Fe-4S] cluster shifted the oligomerization state toward monomers. This observation is consistent with findings for *Ec*HemW, in which the incorporation of the iron-sulfur cluster likewise promotes dimer formation.^61^

In contrast to *Ec*HemW, which has been reported to bind heme covalently, our spectroscopic and titration assays demonstrated that *Mc*HemW binds heme noncovalently. Given that free heme concentrations in the cell are maintained at extremely low levels, heme binding by *Mc*HemW *in vivo* likely requires facilitation through protein-protein interactions from the other heme donors, and this process could also be dominated by the thermodynamics of the affinity difference between heme carriers. Although a specific heme-binding site in HemW enzymes has not yet been experimentally confirmed, the conserved HNX_2_YW motif uniquely present in HemW has been proposed as a putative heme-binding region.^62,96^ For comparison, cytochrome c family members employ a CX_2_CH motif in which the two cysteines form covalent, thioether bonds with the heme vinyl groups, anchoring the cofactor to the protein backbone, while the histidine often serves as an axial ligand to the heme iron.^97^

Among HemN-like radical SAM enzymes involved in heme biology, ChuW, HutW, and HmuW are classified as class C radical SAM methyltransferases that function in the anaerobic heme degradation pathway.^50,54,98–100^ These enzymes are proposed to use two equivalents of SAM, one to generate a 5’-dAdo radical, and the other to undergo homolytic C-S bond cleavage, producing anaerobilin and related analogs as end products.^47,48,50,98,101^ In these systems, 5’-dAdo and SAH are generated in the absence or the presence of heme.^40,54^ Other known members of the radical SAM enzyme superfamily that utilize two equivalents of SAM are those that catalyze radical-based one-carbon transfer reactions to install methyl groups or cyclopropane moieties in a variety of natural products.^30,43,45,102^ In our SSN analysis, ChuW, HutW, and HmuW consistently formed a tight cluster but did not group with *Mc*HemW, indicating a more distant evolutionary relationship (Figure S2). This divergence may explain the weak SAM cleavage activity we observed for *Mc*HemW compared to *bona fide* heme-degrading enzymes. The low residual enzyme activity observed for *Mc*HemW was enhanced in the presence of heme and was not biased toward either 5’-dAdo or SAH production, suggesting that heme may play a role in steps preceding 5’-dAdo radical production. It has been proposed that the heme-Fe center could facilitate electron transfer in HutW when NADPH is used as the reductant.^51^ The same mechanism may explain the increased activity in heme-reconstituted *Mc*HemW. Nonetheless, the influence of heme on the activity at the RS site suggests a plausible interaction between heme and the [4Fe-4S] cluster in *Mc*HemW.

## Conclusion

Using bioinformatics, biochemical, and spectroscopic approaches, we characterized the *hemW* gene in *M. catarrhalis* as a homolog of *Ec*HemW within the radical SAM superfamily. Sequence similarity network analysis placed *Mc*HemW in close evolutionary proximity to *Ec*HemW, a putative heme chaperone. Biochemical assays supported the presence of a [4Fe-4S] cluster that is essential for dimerization. Spectroscopic and mass spectrometry analyses demonstrated specific, noncovalent heme binding, while activity assays revealed that heme enhances *Mc*HemW-mediated radical-based reaction. Our *in vitro* characterization of *Mc*HemW provides insights that will facilitate functional investigation of *Mc*HemW and the exploration of its biological role as a previously unrecognized contributor to iron homeostasis in this human pathogen.

## Notes

The authors declare no competing financial interest.

## Acknowledgments

The authors thank Dr. Anthony T. Iavarone at the University of California, QB3 Mass Spectrometry Facility, Dr. Huan He at the FSU College of Medicine Translational Science Laboratory, and Dr. Xingsong Lin at the MASS Lab in the Department of Chemistry and Biochemistry for support with protein mass spectrometry analysis. We also thank Dr. Christopher Lemon at Montana State University for the insightful discussion. This work was supported in part by the National Institute of General Medical Sciences of the National Institutes of Health under Award Numbers R35GM160337 to W.Z., R35GM126961 to R.D.B., and R35GM156215 to VdC-L; the content is solely the responsibility of the authors and does not necessarily represent the official views of the National Institutes of Health. The work was also partially supported by the Florida State University Start-up Fund to W.Z., the National Science Foundation CHE-2320338 to W.Z., and the Katherine Blood Hoffman Endowed Scholarship offered by the Florida State University Department of Chemistry and Biochemistry to C.P.

## TOC

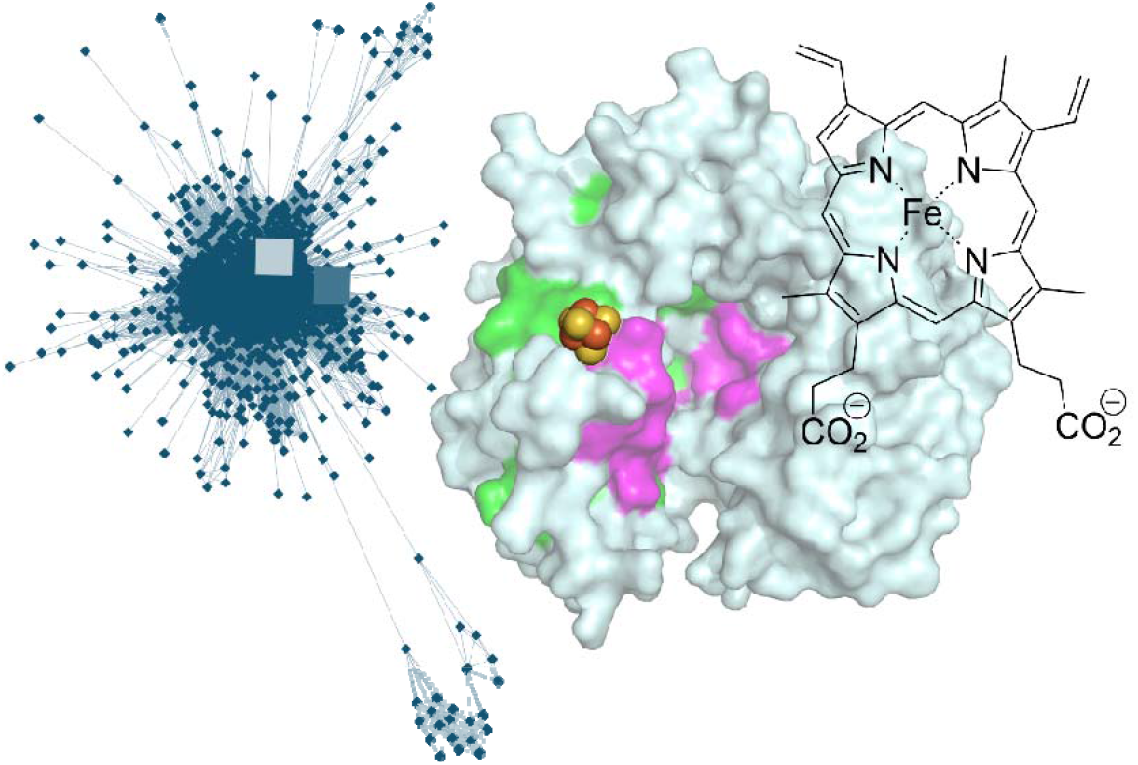

