## SupplementaryInfo for "*Moraxella catarrhalis* HemW is a Heme-binding Radical SAM Enzyme"

**Materials and Methods**

**General information.** All chemicals and reagents were purchased from Merck Sigma-Aldrich and Fisher Scientific unless otherwise noted. Plasmid DNA was synthesized by GenScript (Piscataway, NJ). DNA sequencing was performed at the Biological Core Facility at Florida State University. Protein mass spectrometry (MS) analysis was performed at the QB3 Mass Spectrometry Facility, University of California, Berkeley (Berkeley, CA). Amino acid analysis was performed at the Molecular Structure Facility at the University of California, Davis. Primers used for site-directed mutagenesis were obtained from Eurofins (Louisville, KY). Anaerobic experiments were conducted in a mBraun Unilab Pro SP anaerobic chamber equipped with an O_2_ sensor (Stratham, NH).

**Bioinformatic analysis.** Sequence similarity analysis was performed using the RadicalSAM.org Sequence Similarity Network (SSN), a curated database of networks of radical *S*-adenosylmethionine (SAM) enzymes based on sequence homology. *Mc*HemW was queried to determine its cluster membership and relatedness to other radical SAM enzymes. The SSN was examined across a range of alignment score thresholds to evaluate how *Mc*HemW grouped with related sequences at varying levels of sequence similarity. Additional sequences listed in Table S1 were similarly queried using their respective UniProt accession numbers to identify their cluster locations and subcluster assignments within the SSN. Multiple sequence alignment was performed using MAFFT, and the resulting alignments were visualized using Jalview to assess conservation of functionally relevant motifs. To analyze the gene neighborhood of *hemW* in bacteria, we first searched for HemW homologs in diverse bacterial phyla via BLAST. Homologs were confirmed on the basis of containing the HemW signature motif HNX_2_YW and a typical three-cysteine radical SAM motif CX_3_CX_2_C, as opposed to the four-cysteine motif CX_3_CX_2_CX_2_C found in HemN. Gene neighborhoods from the diverse bacteria were initially analyzed using *fast.genomics*^1^ and conserved gene neighborhoods were then visualized using Gene Graphics^2^.

**Construct design and protein expression.** The gene encoding the *M. catarrhalis* HemW protein was codon-optimized for *E. coli* expression, synthesized, and inserted into the pET28b vector between the *NdeI* and *BamHI* restriction sites, resulting in the N-terminal His-tagged *Mc*HemW plasmid. All the *Mc*HemW variants were generated using the QuikChange method with the primers listed in Table S2. DNA sequences of these mutations were confirmed by Sanger DNA sequencing using T7 forward and reverse primers. The plasmid encoding the *Mc*HemW gene was transformed into *E. coli* TOP10 competent cells for plasmid amplification and BL21-Gold (DE3) competent cells for protein expression. The expression condition was adapted from the previously published protocol with modifications. Briefly, the transformed *Mc*HemW-BL21-Gold (DE3) cells were grown overnight in LB broth containing 50 μg/mL kanamycin at 37 °C. 10 mL of the overnight culture was used to inoculate 1 L of LB broth containing 50 μg/mL kanamycin. The culture grew at 37 °C with shaking at 220 rpm until the optical density at 600 nm (OD_600_) reached approximately 0.6. The incubation temperature was then reduced to 18 °C with shaking at 90 rpm for 1 hour. Protein expression was induced after the 1-hour cooling period by the addition of isopropyl-β-D-thiogalactopyranoside (IPTG) to a final concentration of 0.3 mM. At the time of induction, cysteine, magnesium sulfate, and ammonium iron(III) citrate were also added to final concentrations of 1.5 mM, 0.75 mM, and 0.15 mM, respectively. The culture grew at 18 °C with shaking at 90 rpm for an additional 18 hours. Cells were harvested by centrifugation at 4,000 rpm for 20 minutes at 4 °C, and the resulting cell pellet was frozen and stored in liquid nitrogen before purification.

**Protein purification.** All purification steps were conducted in the mBruan anaerobic chamber with O_2_ maintained below 1 ppm. The purification method was adapted from the previous work with modifications.^3,4^ The frozen cell pellet was resuspended in Buffer A (50 mM Tris-Cl, 150 mM NaCl, 30 mM imidazole, pH 7.5), with the addition of 1 mM dithiothreitol (DTT), 1x BugBuster®, 100 µM ammonia Fe(III) citrate, 10 U/mL benzonase, and 0.5 mg/mL lysozyme with continuous stirring for 1 hour. The cell lysate was transferred to sealed centrifuge tubes and centrifuged at 12,000 rpm for 45 minutes at 4 °C outside of the anaerobic chamber. After centrifugation, the supernatant was transferred to the anaerobic chamber and loaded onto a Ni-NTA column pre-equilibrated with Buffer A. The column was washed with at least five column volumes of Buffer A containing 1 mM DTT. *Mc*HemW was eluted with Buffer B (50 mM Tris-Cl, 150 mM NaCl, 300 mM imidazole, 1 mM DTT, pH 7.5). The colored elution was collected and concentrated using an Amicon Ultra 0.5 ml 30K spin filter, with each concentration step performed for 12 minutes at 14,000 x g. The concentrated protein was loaded onto a PD-10 column equilibrated with Buffer C (50 mM Tris-Cl, 150 mM NaCl, pH 7.5) to remove imidazole. When chemical reconstitution was performed, ammonium Fe(III) citrate and sodium sulfide were added to the protein sample after the initial PD-10 desalting step and incubated at room temperature while mixing for 30 minutes. The sample was then passed through a second PD-10 column equilibrated with Buffer C to remove excess iron and sulfur. Protein samples were further concentrated before being flash-frozen and stored in liquid nitrogen immediately after removal from the anaerobic chamber. Protein concentration was determined using a Bradford assay with bovine serum albumin as the standard. A calibration curve was generated by measuring absorbance at 595 nm after incubation with Bradford reagent. The measured HemW protein concentration was corrected by a factor of 0.65, determined by amino acid analysis.

**Iron content determination.** The iron content assay was performed under aerobic conditions using a N,N-dimethyl-*p*-phenylenediamine dihydrochloride (ferrozine)-based method, as previously reported.^5^ The assay required the preparation of the following solutions: 22% (v/v) HNO_3_, 7.5% (w/v) ammonium acetate, freshly prepared 12.5% (w/v) ascorbic acid, and 10 mM ferrozine. A standard curve was generated using iron standards diluted with water at concentrations ranging from 0 µM to 1000 µM. For each standard or sample, 100 µL was incubated in triplicate with 100 µL of 22% HNO_3_ at 90 °C for 30 minutes. After incubation, samples and standards were cooled to room temperature and centrifuged at 15,000 x g for 5 minutes to pellet any precipitate. 600 µL of 7.5% ammonium acetate was added to neutralize the solution. Subsequently, 100 µL of 12.5% ascorbic acid and 100 µL of 10 mM ferrozine were added to each tube. The mixtures were centrifuged again at 15,000 x g for 5 minutes, and the supernatants were transferred to cuvettes. Absorbance at 562 nm was measured using a UV-Vis spectrophotometer. A linear standard curve was plotted to determine the iron content in the samples.

**Sulfur content determination.** The sulfur content assay was performed under anaerobic conditions following an established method.^5,6^ The following solutions were prepared: 2 mM sodium sulfide stock (Na_2_S·9H_2_O), 1% (w/v) zinc acetate, 12% (v/v) NaOH, 0.1% (w/v) ferrozine, and 10 mM FeCl_3_. A sulfide standard curve was generated using dilutions of the sodium sulfide stock to achieve concentrations ranging from 0 µM to 1000 µM. For each standard or sample, 200 µL was incubated in triplicate with 600 µL of 1% zinc acetate and 50 µL of 12% NaOH for 15 minutes at room temperature. The samples and standards were centrifuged at 15,000 × g for 1 minute. After centrifugation, 150 µL of 0.1% ferrozine and 150 µL of 10 mM FeCl_3_ were added to each tube. The mixtures were mixed thoroughly and incubated at room temperature for 30 minutes. Tubes were then removed from the anaerobic environment, and absorbance at 670 nm was measured using a UV-Vis spectrophotometer. A linear standard curve was plotted to determine the sulfide content in the samples.

**UV-vis absorption spectroscopy.** Freshly purified protein samples were diluted to 10 µM in the anaerobic chamber, and the resulting solutions were transferred to 1 mL quartz cuvettes sealed with screw caps before being removed from the chamber. UV-vis spectra were recorded using an Agilent Cary 60 spectrometer. For DTH-treated samples, 250 µM of DTH was added to 10 µM of the protein solution in the anaerobic chamber, followed by a 10-minute incubation before being transferred to the cuvette for measurement.

**Anaerobic size exclusion chromatography.** The oligomerization state of *Mc*HemW was determined using fast protein liquid chromatography (FPLC) with a Cytiva Superdex 200 Increase size-exclusion chromatography column. *Mc*HemW samples at various concentrations (10, 30, 40, 50, 100, and 200 µM) were injected into the column, which had been equilibrated with degassed Buffer C. Elution was performed using the same buffer at a flow rate of 0.22 mL/min, and the absorbance at 280 nm was monitored. This was repeated with 50 µM of the *Mc*HemW ∆RS variant. Commercially available protein standard (Sigma Protein Standard Mix 15-600 kDa) was run on the same column under identical conditions to generate a standard curve. The molecular weight of *Mc*HemW was estimated based on its elution time relative to the standard curve. To assess the monomer-dimer equilibrium, peak areas corresponding to the monomer and dimer species were integrated using Cytiva UNICORN 7.10 evaluation software.

**Electron paramagnetic resonance (EPR) spectroscopy.**

*Sample preparation.* All EPR samples were prepared in an anaerobic chamber (Coy Laboratories, 98% N_2_, 2% H_2_). Glycerol was added to a final concentration of 15% in the 200-300 µM *Mc*HemW sample in Buffer C for all EPR measurements. For dithionite (DTH)-reduced samples, 10 equivalents of DTH were added to the *Mc*HemW sample, which was then transferred to an X-band EPR quartz tube and incubated for 10 minutes in the anaerobic chamber before being flash-frozen in liquid nitrogen inside the chamber. For the sample containing K^13^CN, about 100 equivalents of K^13^CN was mixed with the reduced *Mc*HemW for 2 minutes and was then flash frozen in liquid nitrogen.

*Continuous wave (CW) EPR experiments.* CW EPR measurements were carried out at X-band (~9.4 GHz) using a Bruker EleXsys E500 spectrometer equipped with a superhigh Q resonator (ER4122SHQE). Cryogenic temperatures were maintained using an Oxford Instruments ESR900 continuous-flow liquid helium cryostat with a temperature controller (Oxford Instrument ITC503). Spectra were collected at 10 K unless otherwise noted. Spectrometer parameters were as follows: 60 ms conversion time, 0.2 mW power, 5 G modulation amplitude, and 100 kHz modulation frequency. Spectral simulations were performed using EasySpin 6.0.4 in Matlab.

*Pulsed EPR experiments.* Hyperfine sublevel correlation (HYSCORE) spectra were obtained at X-band and 10 K on a Bruker Biospin EleXsys E-580 spectrometer using a MS5 split-ring resonator. The standard four-pulsed sequence for HYSCORE π/2−τ–π/2–t1−π–t2−π/2−τ–echo with 8-step phase cycling was programmed in PulseSPEL via the XEPR interface. A pulse-length of 12 ns was used for the π/2 pulse (*t*_π/2_) and 24 ns for the inversion pulse (*t*_π_). Time domain spectra were baseline-corrected (third-order polynomial), apodized with a hamming window, zero-filled to 8-fold points, and fast Fourier-transformed to yield the frequency domain spectra. Spectrometer settings were: microwave frequency of 9.195 GHz, magnetic field of 338.6 mT, and τ of 140 ns for *g* = 1.94; The time increment in both dimensions was 24 ns with 180 steps.

**Heme binding assays.** Heme reconstitution was performed under anaerobic conditions. A heme stock solution was prepared by dissolving 4 mg of heme in 400 µL of 50 mM NaOH. The solution was mixed and incubated for 30 minutes at room temperature to ensure complete dissolution. Following incubation, 400 µL of 1 M Tris-Cl was added to the solution, which was then centrifuged at 15,000 x g for 12 minutes. The concentration of the resulting supernatant was determined to be 2 mM based on the extinction coefficient of 58.4 mM^-1^cm^-1^ at 385 nm.^7^ 10 µM of heme solution was added to 10 µM *Mc*HemW and incubated overnight at 25 °C for 16 hours. Binding controls include 10 µM BSA with and without 10 µM of heme. Reduced scan upon heme binding includes 250 µM DTH, a 10-minute incubation with 10 µM protein in the presence or absence of 10 µM heme. Heme titration experiments were performed in the same manner with 0, 0.5, 1, 3.5, 5, 10, and 15 µM heme added to 5 µM *Mc*HemW. The same titration experiment was performed in the absence of protein. All samples were transferred to sealed cuvettes and analyzed using a UV-Vis spectrometer.

**Heme staining PAGE assays**. To assess heme binding by SDS-PAGE and native-PAGE, we analyzed WT *Mc*HemW and heme-reconstituted *Mc*HemW samples alongside the cluster-knockout variant and appropriate controls, including cytochrome c, hemoglobin, and free heme. An equal molar amount of heme solution was added to McHemW and incubated overnight at 25 °C for 16 hours prior to measurement. The same denatured marker, Precision Plus Protein Dual Color Standard, was used for SDS-PAGE and native-PAGE. All samples were run on Mini-PROTEAN TGX gels under standard electrophoresis conditions. The heme staining solution containing 3,3′,5,5′ tetramethylbenzidine (TMBZ) and sodium acetate was freshly prepared in the dark. 6.3 mM TMBZ dissolved in methanol and mixed with 0.25 M sodium acetate buffer, pH 5.0, in a 7:3 ratio. Following electrophoresis, the gel was incubated in the staining solution in the dark for 1.5 hours with occasional mixing. Incubation was followed by the addition of 30% H_2_O_2_ to a final concentration of 0.3% (V/V), and heme staining became visible within approximately 30 minutes.^8^ After the TMBZ reaction, the same gels were stained with Coomassie Brilliant Blue to visualize the protein.

**HPLC-based enzyme activity determination.** All enzymatic activity was performed at room temperature under anaerobic conditions. The enzymatic reaction was initiated by adding 50 µM *Mc*HemW to a reaction mixture containing 2 mM SAM and 1 mM DTH. Reactions were quenched by the addition of 0.5 M formic acid before centrifuging at 14,800 rpm for 10 minutes. The supernatant was subjected to 80°C for 5 minutes before being centrifuged again at 14,800 rpm for an additional 10 minutes. The resulting supernatants were analyzed using a Thermo Vanquish HPLC equipped with a Thermo Scientific Hypersil Gold C18 column using a linear gradient elution method. The mobile phase consisted of A: 1 mM ammonium acetate, pH 5.0, and B: acetonitrile. The flow rate was maintained at 0.25 mL/min, and the absorption at 260 nm was monitored. The gradient began at 0% acetonitrile and increased to 15% over 10 minutes, followed by an increase to 80% B at 25 minutes. The percentage of solvent B was then increased to 100% at 25 minutes and was held at 100% B until 30 minutes, after which the gradient was returned to 0% B by 35 minutes. The column was re-equilibrated at 0% acetonitrile until the end of the 40-minute run. An injection volume of 10 μL was used for both standards and samples. Authentic standards, including SAM, *S*-adenosylhomocysteine (SAH), and 5’-dAdo, were prepared at a concentration of 100 µM and subjected to the same HPLC method. Peaks corresponding to reaction products and standards were identified based on their retention times. Various concentrations of SAH and 5’-dAdo (1, 5, 10, 20, 40, 80, and 100 µM) were ran using the same method to form a calibration curve for the product. The concentration of the product is quantified by the peak area corresponding to the standards with known concentration. Data acquisition and processing were performed using Thermo Chromeleon software.

**LC-MS analysis.** The same reaction mixtures and standards used for activity determination by HPLC were also used for LC/MS analysis. The reaction mixture was injected into the Vanquish UHPLC system coupled to the Thermo TSQ Quantis Mass Spectrometer (LC-MS) system, equipped with a Thermo Scientific Hypersil Gold C18 Selectivity column. The gradient elution used 1 mM ammonium acetate, pH 5.0, and acetonitrile, both containing 0.1% formic acid. The flow rate was maintained at 0.25 mL/min throughout the run. A divert valve was used for 1 minute. After 1 min the gradient began at 0% acetonitrile and increased to 100% over 8.5 minutes and was held at 100% B until 11 minutes. After the gradient was returned to 0% acetonitrile, it was equilibrated for the remainder of the 14-minute run. An injection volume of 10 μL was used. Mass spectra were collected using an electrospray ionization source operated in positive ion mode with Q1 full scan acquisition. The scan range was set from *m/z* 200 to 1000 with a scan speed of 1000 Da per second. The LC flow rate was set to 250 μL per minute. Ion source parameters were as follows: spray voltage 3500 V in positive ion mode and 2500 V in negative ion mode, sheath gas 50 arbitrary units, auxiliary gas 10 arbitrary units, sweep gas 1.0 arbitrary units, ion transfer tube temperature 325°C, and vaporizer temperature 350°C. Data acquisition and processing were performed using Thermo Xcalibur and FreeStyle software.

**Table S1. Protein sequences used in the sequence alignment.**

| **Name** | **Organism** | **Uniprot Code** |
| --- | --- | --- |
| *Mc*HemW | *Moraxella catarrhalis* | A0A3A9RUY7 |
| *Ec*HemW | *Escherichia coli (strain K12)* | P52062 |
| *El*HemW | *Enterobacter ludwigii* | G8LDY9 |
| *Kp*HemW | *Klebsiella pneumoniae* | A6TDW5 |
| *Mo*HemW | *Moraxella ovis* | A0A378PLG8 |
| *Ab*HemW | *Acinetobacter baumannii* | Q6FEY5 |
| ChuW | *Escherichia coli O157:H7* | A0A384LP51 |
| HutW | *Vibrio alginolyticus* | A0A068BDD1 |
| HemZ | *Bacillus subtilis (strain 168)* | Q796V8 |
| *Ec*HemN | *Escherichia coli (strain K12)* | P32131 |

**Table S2. DNA and protein sequences of *Mc*HemW and primer sequences used for site-directed mutagenesis**

| **Name** | **Sequence** |
| --- | --- |
| C22A-fwd primer | 5’-ATCCACATTCCGTGGGCTGTTAAAAAGTGCCCG-3’ |
| C22A-rev primer | 5’-CGGGCACTTTTTAACAGCCCACGGAATGTGGAT-3’ |
| C29A-fwd primer | 5’-AAAAAGTGCCCGTATGCCGACTTCAACAGCCAT-3’ |
| C29A-rev primer | 5’-ATGGCTGTTGAAGTCGGCATACGGGCACTTTTT-3’ |
| *Mc*HemW DNA sequence (codon-optimized) | ATGTATCCCGACCTATTAGATCCAGCTCACATCCCGCTCAGCCTGTATATCCACATTCCGTGGTGTGTTAAAAAGTGCCCGTATTGCGACTTCAACAGCCATGCACTGCCTAGTCAAGTTCCGTTCGAGAACTACATCGAGGCGTTGCTGAGCGACGCTGTGAGCCAACAAAGCCTCGTATCCAACCGTCAGATTGACACCGTTTTTATCGGCGGTGGTACACCGTCGCTGCTGCCGATTGAAAGCTTTAAACGTTTGTTTGACGGCTTGCGCTTAATTTATGATTTCGCCCCGACGTGCGAAATTACCTTGGAGGCTAATCCGGGTACGTTGGAACACAGCCCGTTCGACGAATATCTGACTATTGGGATCAACCGCCTGTCGATCGGTGTCCAGTCTTTCGACCACAACGCTTTGACCGTGCTGGGCCGTATCCACAACCCGGACCAGGCAAGCAATGCCATTCGTGCCGCACGTATTGCTGGCTTCGACCGCGTGAACGCCGATCTGATGTACGGCCTGCCGAACCAGAATCCGAGCAAAGCGGTTAACGACATCCAGACCGCGATAGATGCGGGTGCGACCCATCTGTCCTGGTATCAACTGACTATTGAGCCGAATACCACCTTTTACTGCACCCCGCCTCAGCTGCCGGACGAGGAGATCATGGCGGAAATCGAAACCCTGGGTCAGTCCGTGCTGACCAAGCACGGCTTCTACAATTACGAAGTTAGCGCGTGGCATGGTCCGAATGACGCGCCGTGTCAGCACAATCTGAATTACTGGCAGTTTGGTGATTATCTGGCAATCGGCGCAGGCGCGCATGGTAAACTGACTCTTAAGAACCATCCGGACTACCAAGATGGCATTTACCGCTACCAAAAATCTCGTCAGCCGAAAGACTATTTGGCGTTCGAGGATGCTCCGAAGATGGTTAACTTTGAACGTATTTCTGACGATAACTTACCGTTCGAGTTCATGATGAATGCGCTGCGTCTGAAGGATGGTGTTTCGACCGATGTCTTCGAGACGCGTACCGGTTTGCTGATTAGCACGATCGATAACGAAATGCTGCCACTTCAACACCAGGGTCTGATGGTGCTGGACCCGCTGCGCATCGCGCCAACCAGACTAGGCTTTCGTTACGTGAACTACCTGGTGTCCCAATTTATCTAA |
| *Mc*HemW protein sequence  (Purification tag originated from the pET28b vector is underlined) | MGSSHHHHHHSSGLVPRGSHMYPDLLDPAHIPLSLYIHIPWCVKKCPYCDFNSHALPSQVPFENYIEALLSDAVSQQSLVSNRQIDTVFIGGGTPSLLPIESFKRLFDGLRLIYDFAPTCEITLEANPGTLEHSPFDEYLTIGINRLSIGVQSFDHNALTVLGRIHNPDQASNAIRAARIAGFDRVNADLMYGLPNQNPSKAVNDIQTAIDAGATHLSWYQLTIEPNTTFYCTPPQLPDEEIMAEIETLGQSVLTKHGFYNYEVSAWHGPNDAPCQHNLNYWQFGDYLAIGAGAHGKLTLKNHPDYQDGIYRYQKSRQPKDYLAFEDAPKMVNFERISDDNLPFEFMMNALRLKDGVSTDVFETRTGLLISTIDNEMLPLQHQGLMVLDPLRIAPTRLGFRYVNYLVSQFI |

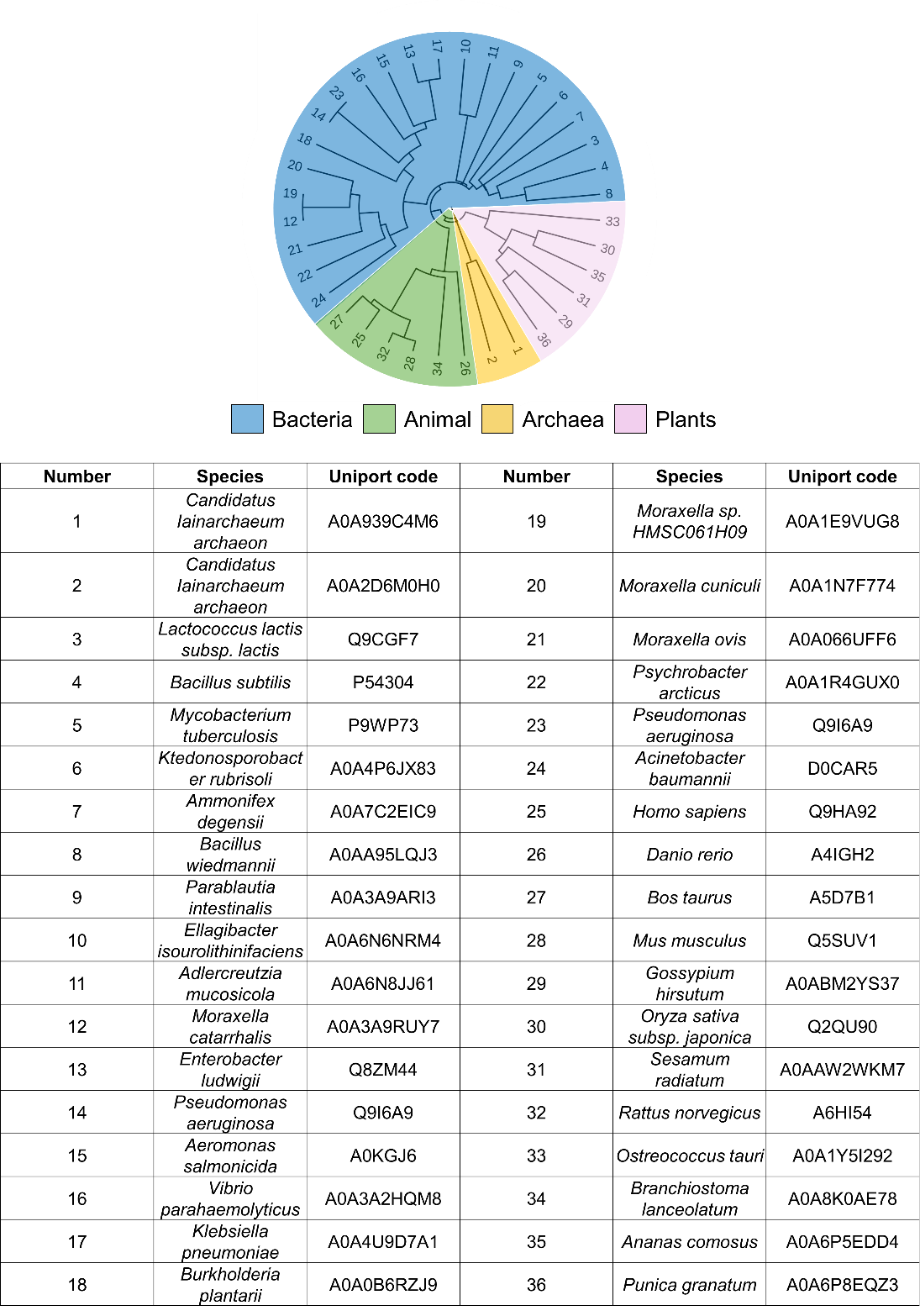

**Figure S1. HemW across all kingdoms.** Phylogenetic tree showing the species relationships of HemW throughout bacteria (blue), animals (green), archaea (yellow), and plants (pink). Table lists species and the uniport code used for sequence alignment.

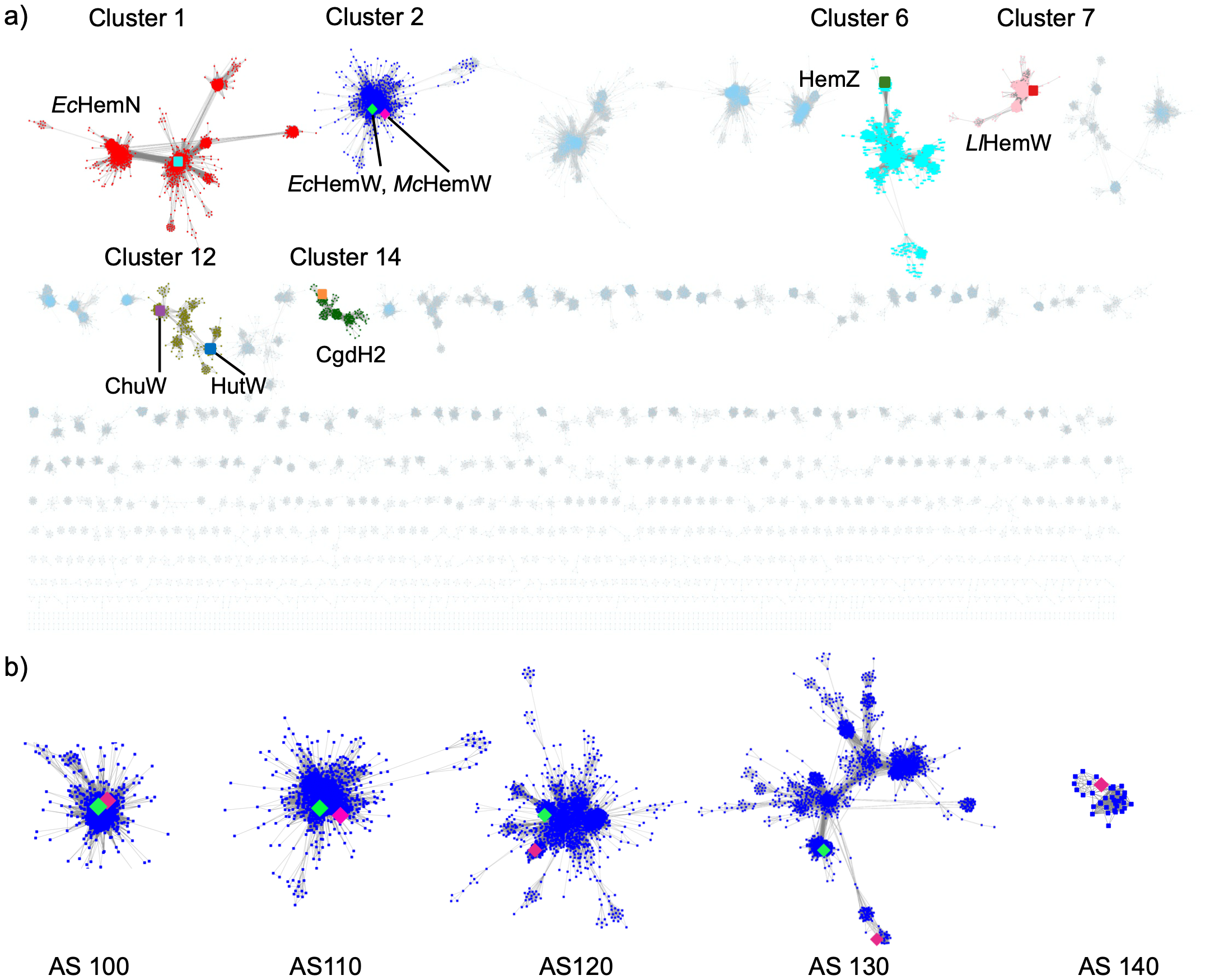
 **Figure S2. Megacluster 2-2 oxygen-independent coproporphyrinogen III oxidase-like enzymes.** a) Sequence similarity network (SSN) of RadicalSAM.org Cluster 2-2 at alignment score 110, with clusters of interest highlighted. Cluster 1 (red) contains *Ec*HemN (cyan). Cluster 2 (dark blue) includes *Ec*HemW (green) and *Mc*HemW (pink). Cluster 6 (cyan) contains HemZ (dark green), Cluster 7 (pink) contains *Ll*HemW (red), Cluster 12 (tan) includes ChuW (purple) and HutW (blue), and Cluster 14 (green) contains *Campylobacter jejuni* CgdH2 (orange). All nodes are connected by gray edges representing sequence similarity, while clusters not of interest are shown in light blue. b) Clusters containing *Mc*HemW throughout various alignment scores of 100-140. Highlighted in green is *Ec*HemW if present in the cluster and *Mc*HemW in pink.

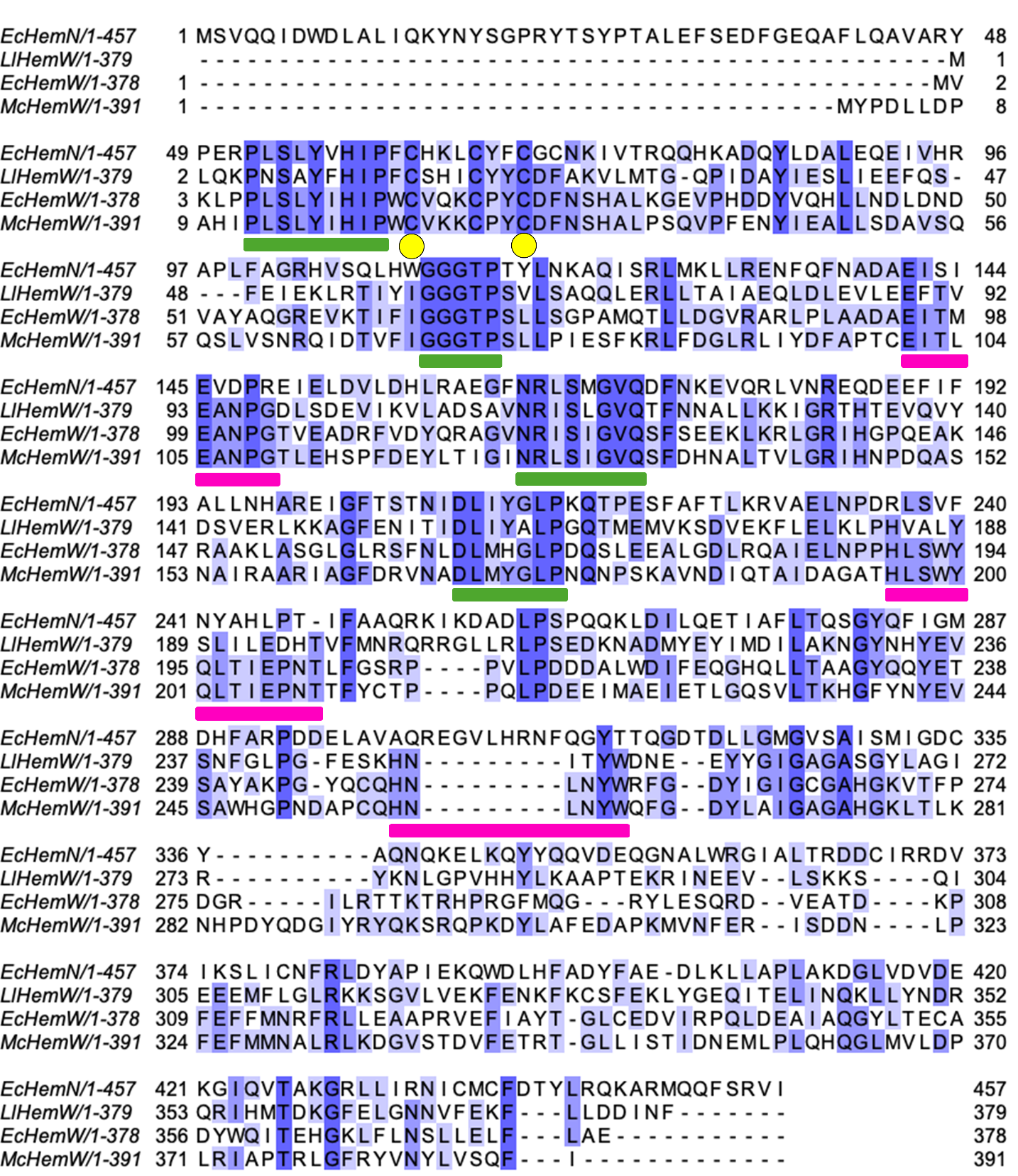

**Figure S3.** **Multiple sequence alignment of representative HemW and HemN**. The alignment includes *Ec*HemN, *Ll*HemW, *Ec*HemW, and *Mc*HemW, generated using Clustal Omega and visualized in JalView. Conserved residues are highlighted in blue, and residues targeted for mutagenesis in this study are marked by yellow spheres. Magenta bars denote features conserved across HemW homologs, whereas the green bar marks shared features among heme-binding radical SAM enzymes such as HemN.

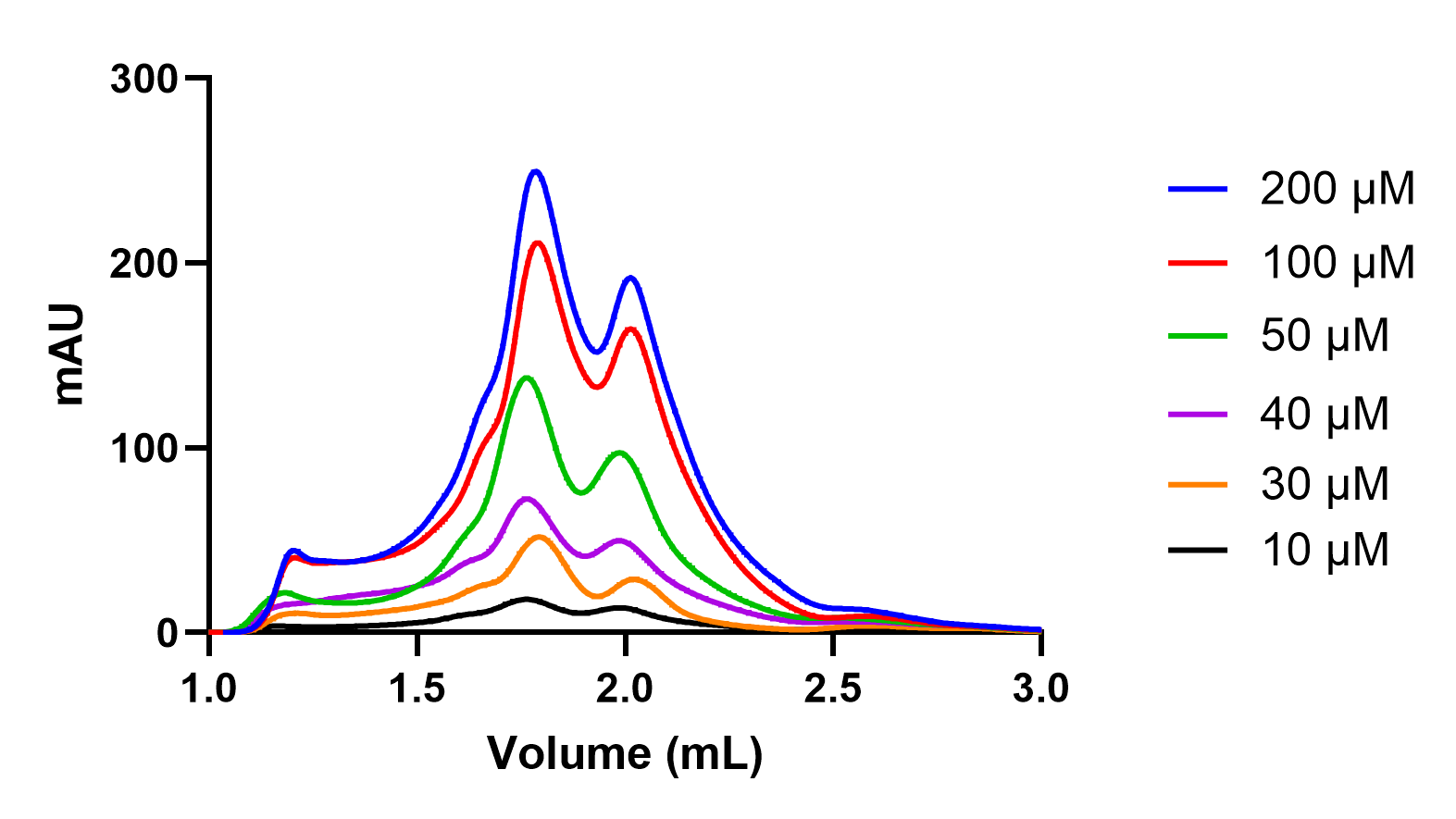

**Figure S4. Size-exclusion chromatography profiles of *Mc*HemW at varying protein concentrations.** SEC traces collected at 6 different *Mc*HemW concentrations (10, 30, 40, 50, 100 and 200 µM).

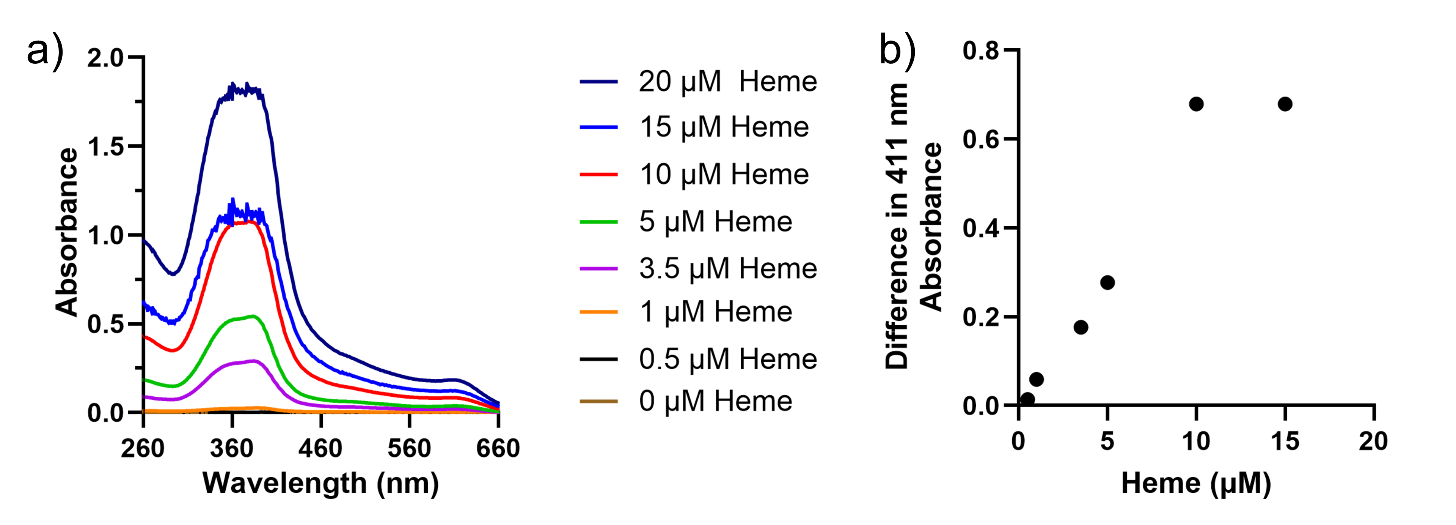

**Figure S5. Free-heme titration standard curve.** a) UV-vis spectrum from 260 nm to 660 nm of heme titration with heme standard in Buffer C with increasing concentrations of heme. b) The difference in absorbance at 411 nm between the heme titration in the presence and absence of *Mc*HemW was calculated from the plot in Figure 5c and Figure S5a. The increasing trend up to 10 µM of heme in the presence of 5 µM of *Mc*HemW, followed by a saturation point, suggests that the elevated 411 nm signal does not simply result from the increased free heme concentration in the background but instead indicates increased protein-bound heme formation. At high heme concentrations, the protein becomes saturated with heme, and no further increase in signal occurs as heme concentration continues to rise.

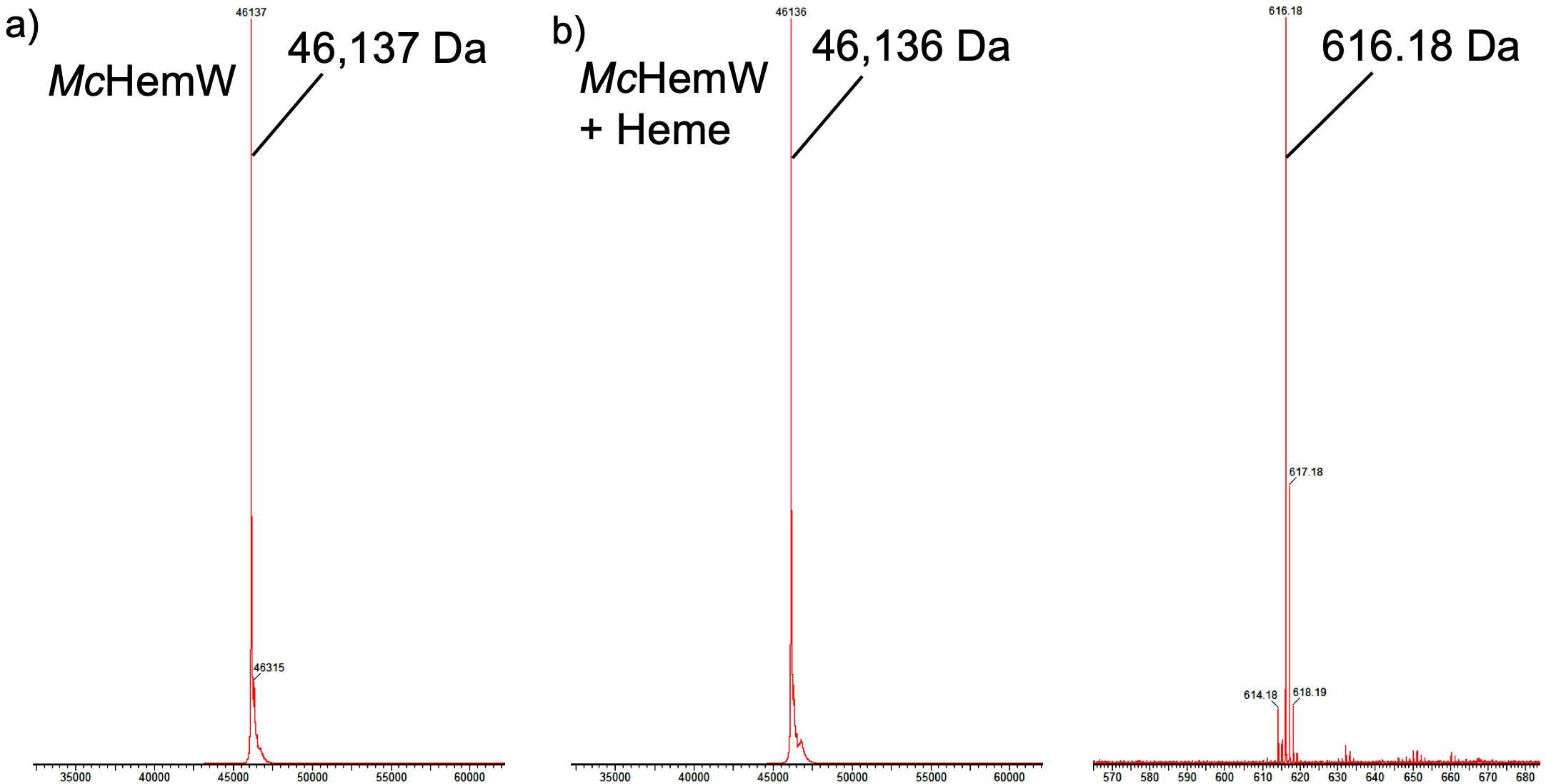

**Figure S6. Intact protein mass spectrometry of as-purified *Mc*HemW and heme-reconstituted sample.** a) Intact mass spectrometry of *Mc*HemW (50 µM) showing a mass of 46,137 Da. b) Spectrum of 50 µM *Mc*HemW after heme reconstitution, showing the expected protein mass of 46,137 Da and a peak at 616.18 m/z corresponding to free heme, with no detectable signal for a heme-HemW adduct.

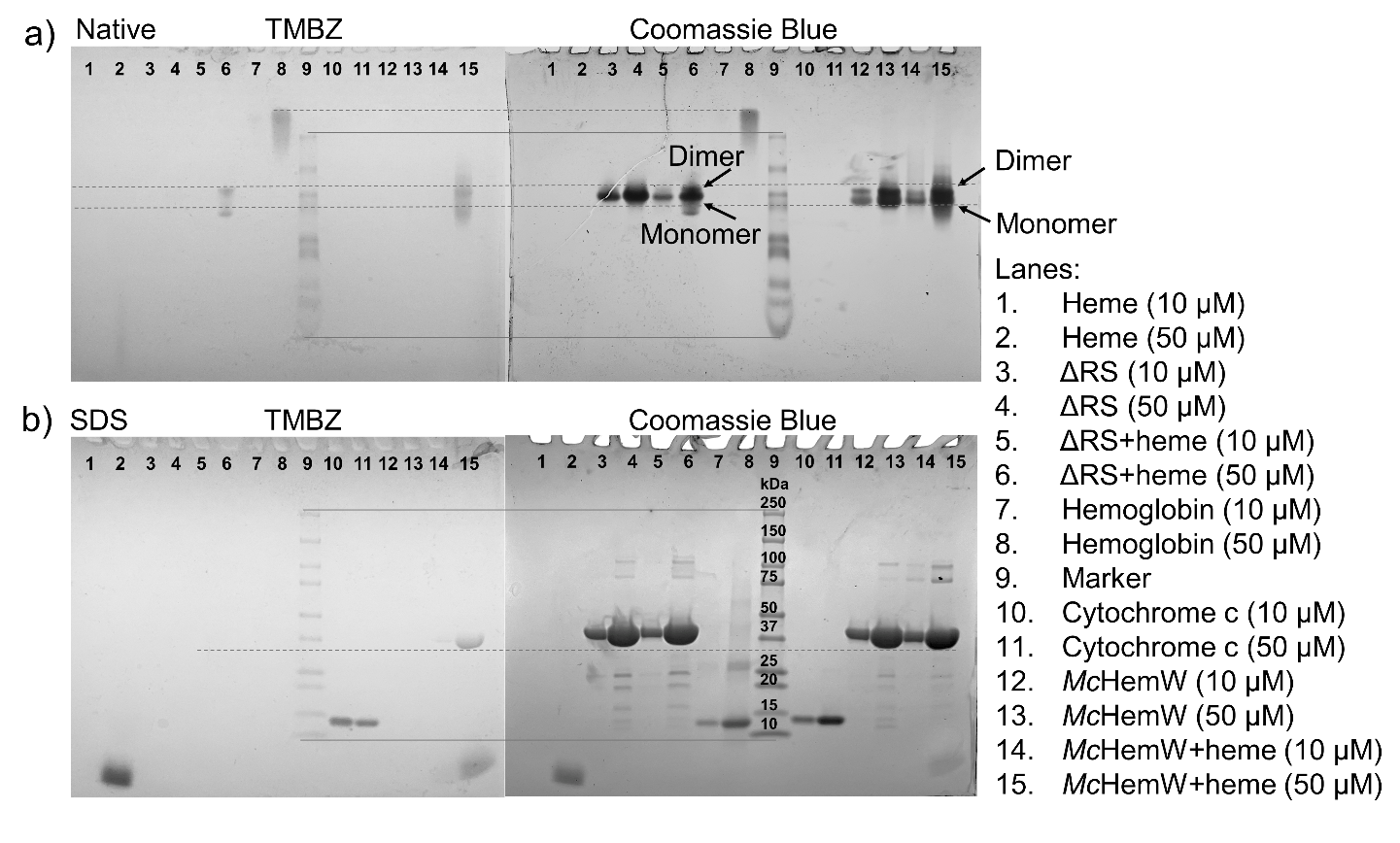

**Figure S7. Native-PAGE and SDS-PAGE analysis of *Mc*HemW and heme-reconstituted *Mc*HemW and ∆RS variant.** a) Native-PAGE analysis of *Mc*HemW visualized using TMBZ heme staining (left) followed by Coomassie Blue staining (right). The protein marker is solely used for alignment. b) SDS-PAGE analysis of *Mc*HemW stained with TMBZ (left) to detect heme-bound species and subsequently with Coomassie Blue (right) to visualize total protein. All gels maintain the same lane order, and the Precision Plus Protein Dual Color Standard was used as the denatured molecular weight marker in both Native PAGE and SDS PAGE.

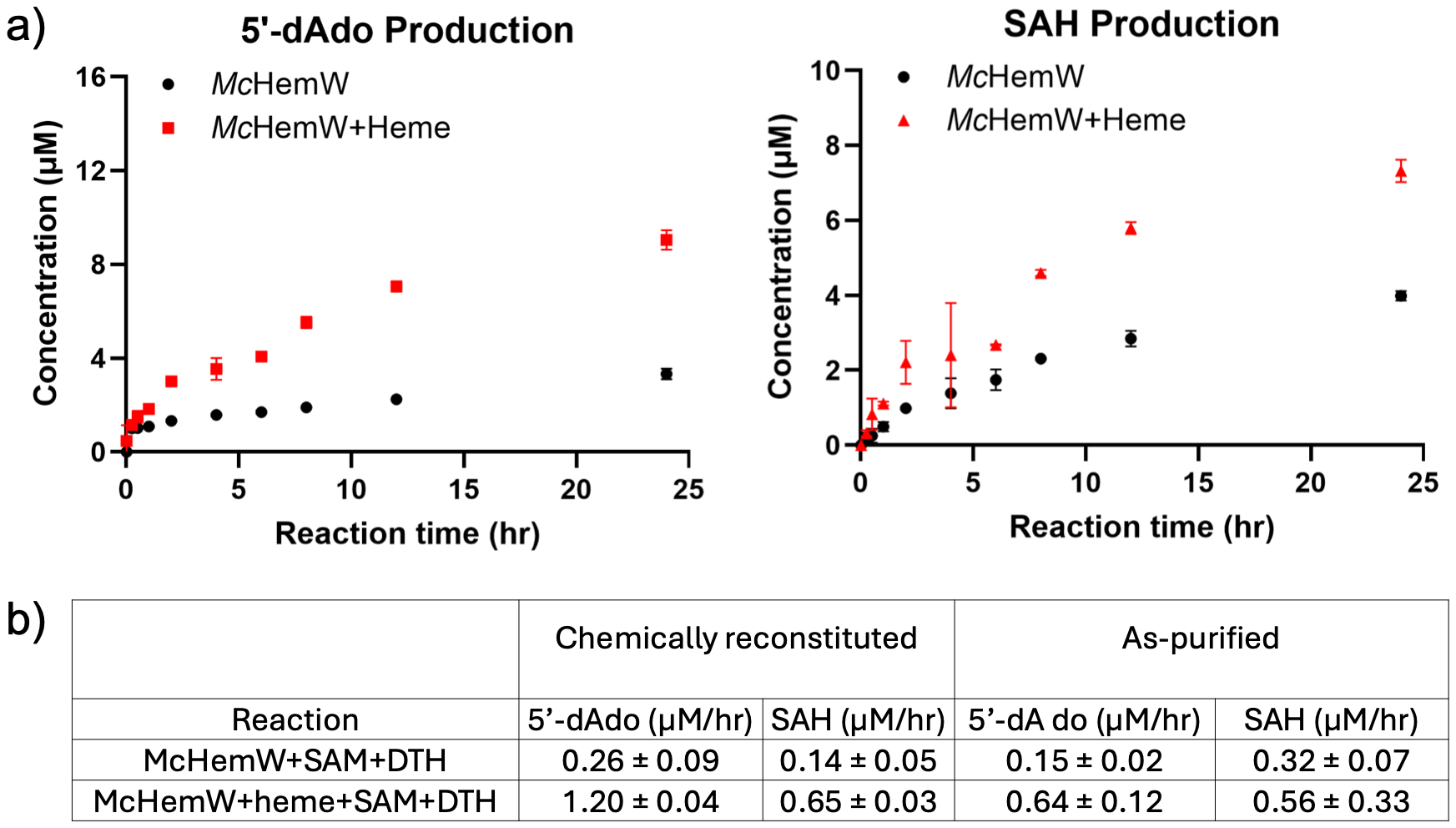
 **Figure S8. Time course of the activity assay of the as-purified *Mc*HemW before and after reconstitution with heme.** a) Time-course analysis of as-purified HemW-catalyzed reactions shows that heme significantly enhances the production of 5’-dAdo and SAH. Error bars represent the standard deviation from two independent experiments. b) Comparison of initial rates of 5’-dAdo and SAH produced from as-purified and chemically reconstituted *Mc*HemW in the presence and absence of heme.

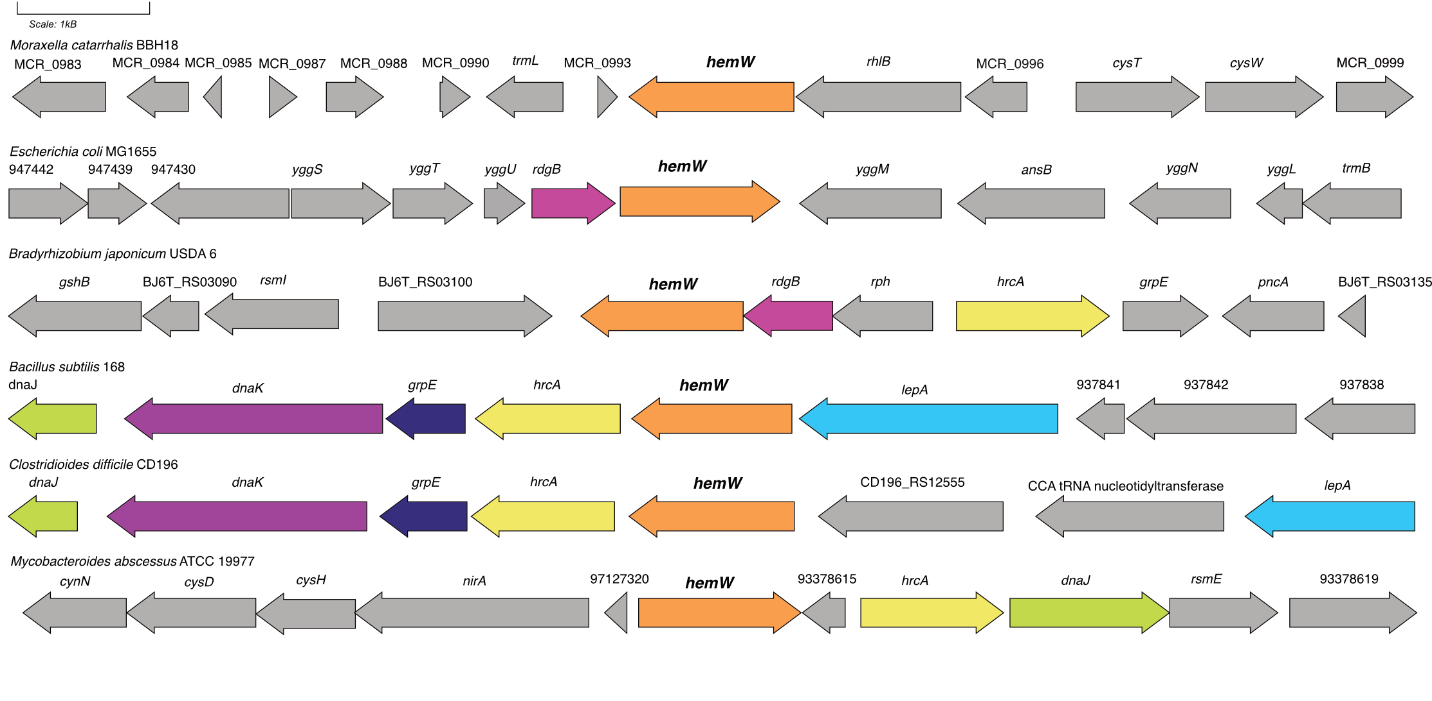

**Figure S9. Gene neighborhoods of *hemW* in diverse Gram-negative and positive bacteria.** Homologous genes that occur more than once are colored, whereas all other genes are in grey. *Moraxella catarrhalis hemW* exists in an operon with the RNA helicase *rhlB* and MCR_0996, a protein of unknown function. In Gram-negative bacteria, such as *E. coli* (Gammaproteobacteria) and *B. japonicum* (Alphaproteobacteria) *hemW* associates with the nucleoside triphosphate-metabolizing pyrophosphatase *rdgB***.** In contrast, Gram-positive bacteria, and select Gram-negative bacteria contain an association of *hemW* with genes related to the heat shock response, including *grpE*, *hrcA*, *dnaK*, and *dnaJ*. This includes *Bacillus subtilis* (Bacilli), *Clostridioides difficile* (Clostridia), and *Mycobacteroides abscessus* (Actinomycetes).
